# Category selectivity observed in the human brain is distinct from category selectivity observed in artificial neural networks

**DOI:** 10.64898/2026.05.29.728609

**Authors:** Alish Dipani, N. Apurva Ratan Murty

## Abstract

Category selectivity for images of faces, scenes, and bodies is among the most striking and reproducible findings in vision neuroscience. Artificial neural networks (ANNs) trained on visual tasks also develop category-selective units, which has led to the suggestion that ANNs may capture important aspects of how the brain processes visual categories. But the mere presence of category-selective units in ANNs does not mean that those units are selective in the same way as the brain. Here, we distinguish between the presence of category-selective units in ANNs from the form of selectivity they express, and show that the selectivity that emerges in ANN units differs in meaningful and systematic ways from that observed in the human brain with fMRI. To this end, we first identified category-selective units in a wide range of ANN models using standard fMRI localizers, and found that selective units emerged reliably in trained, but not in untrained, ANNs. We then identified category-selective regions in the human brain using the same localizer and found that their response tuning to a broad range of images was strikingly consistent across individuals. Thus, category-selective regions exhibit a stable representational signature shared across subjects. Category-selective ANN units did not match this structure. Their responses diverged in both univariate tuning and multivariate representational geometry, fell well below the human-human ceiling, varied substantially across models, and depended strongly on the localizer used to identify them. We also found that the category-selective ANN units were neither necessary nor sufficient for predicting neural responses using an encoding model. Further stimulus-level analyses revealed clear and interpretable mismatches between ANN selectivity and human fMRI responses, which can be used to test and compare better ANN models in the future. Taken together, these results show that the full range of response tuning in category-selective regions provides a highly demanding and discriminative test of brain-model alignment than previously appreciated. Although current ANNs contain category-selective units, the selectivity they express is more fragile and does not capture the stable and shared form of selectivity observed in the human brain.

## 1 Introduction

Few findings in cognitive neuroscience are as robust or influential as category-selective responses in high-level visual cortex. Regions like the fusiform face area (FFA) [1–3], extrastriate body area (EBA) [4, 5], and parahippocampal place area (PPA) [6–8] respond maximally to faces, bodies, and scenes respectively and have shaped how we think about brain organization, development, and evolution [9–23]. Intriguingly, artificial neural networks (ANNs) trained on a wide variety of tasks also develop category-selective units that respond more strongly to faces, bodies, and scenes than to other categories [24–38]. This parallel has led to the suggestion that ANNs may capture the fundamental computational principles underlying visual category selectivity. But this conclusion assumes that the category-selective units in ANNs reflect the same underlying mechanisms and features found in the brain. In this study, we distinguish between the mere presence of category-selective units in ANNs and the form of category selectivity they exhibit, and show that category selectivity that develops in ANN units is different from the selectivity observed in human category-selective

The central challenge in interpreting category selectivity is that the standard definition of selectivity is underconstrained. A voxel in the brain (or unit in an ANN model) is typically deemed, say, face-selective if it responds more to faces than to other categories. But as shown in **Fig 1a**, this minimal criterion admits a wide range of response profiles. For example, both a voxel that responds exclusively to faces and one that responds broadly but slightly more to faces would qualify as face-selective. In fact, there exist *infinitely* many response patterns that can be deemed face-selective (illustrated as the blob in **Fig 1a**). This flexibility in definition invites a subtle but consequential cognitive bias [39, 40]. Once a voxel or unit is labeled “face-selective”, we are inclined to group such units as conceptually similar to the brain, implicitly assuming that they instantiate a common representational principle. But without carefully examining the full range of the response profile and comparing against the profile observed in brains, we cannot know whether observed selectivity in ANNs is truly brain-like.

**Figure 1.**
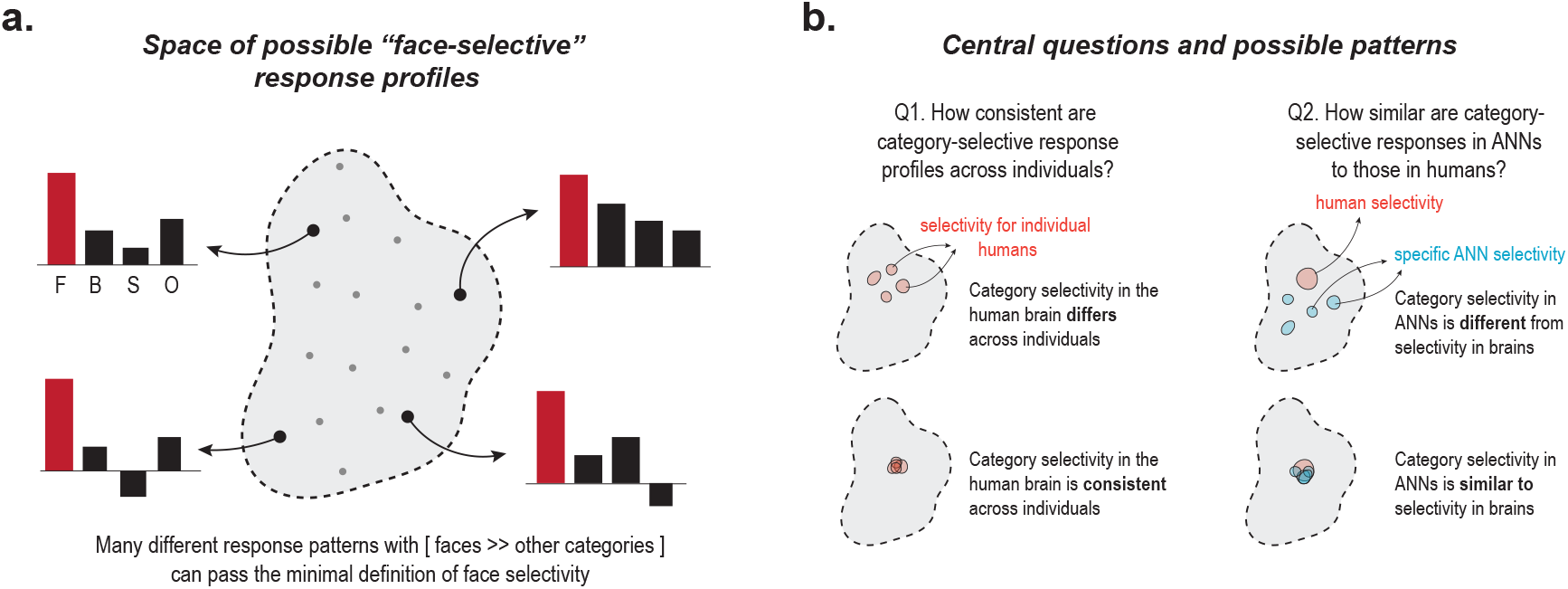
Conceptual framework, questions, and possibility space. **a.** Motivating conceptual framework: The gray blob represents the space of response profiles satisfying the minimal criterion for face selectivity (faces » other categories). Gray and black dots illustrate some sampled face-selective response profiles within the space. Barplots show schematic category-averaged responses for faces (F), bodies (B), scenes (S), and objects (O); faces in red. Many different response profiles can satisfy the minimal definition of face selectivity (faces » other categories). **b.** Schematic illustrating the two central questions explored in this study. Left, possible patterns of category selectivity across humans. Right, possible relationships between category selectivity in humans (red) and ANNs (blue).

This framing leads naturally to two core questions that motivate this study (**Fig. 1b**). First (Q1), how consistent are category-selective response profiles across individuals? If human selectivity patterns vary substantially across people, then the question of comparing any model to a single canonical pattern would itself be ill-posed. Q1 therefore contrasts two possibilities: either selectivity profiles in human category-selective regions are highly idiosyncratic across people, or they are systematic enough to define a stable population-level reference (**Fig 1b, left**). Second (Q2), do ANN units exhibit the same kind of category-selective response profiles as brain voxels in category-selective regions? Q2 thus contrasts two possibilities: One possibility is that brains and ANN units differ in their selectivity profiles despite both being deemed category-selective by the minimal definition. The second possibility is that ANN-units mirror the structure of category selectivity observed in the brain.

To answer these questions, we leveraged 7T fMRI data from the Natural Scenes Dataset [41], which provides stimulus-level responses to thousands of natural images. Using these data, we asked whether category selectivity is consistent across humans and whether category-selective units in ANNs exhibit the same form of selectivity observed in the brain. To foreshadow our results, we found that the pattern of responses from category-selective regions (FFA, EBA, PPA, and others) was highly consistent across humans. This similarity provided a stable target to compare ANN representations. We then applied a localizer-based procedure (like human fMRI) to ANN models and found that face-, body-, and scene-selective units emerge reliably in trained, but not untrained, ANNs. Do these ANN units exhibit the same *form* of selectivity as human category-selective regions? We found that they do not. Although category-selective ANN units preferred the same category of images, their response patterns were reliably different from those seen in human brains with fMRI. These differences were robust to analysis choices and human-interpretable across models and reveal distinct response signatures in brains and ANNs. Together, these findings reveal a dissociation between the existence of category-selective units and the form of category selectivity they exhibit. This distinction shifts the focus from whether category-selective units emerge to how category selectivity is represented, providing a more quantitative framework for comparing biological and artificial visual systems.

## 2 Results

### 2.1 Category-selective units emerge in trained, but not untrained, ANNs

Do artificial neural networks (ANNs) contain face-, body-, and scene-selective units? For a balanced comparison with brains, we subjected several pretrained ANNs to the same localizer images used to identify category-selective regions in the human brain [42–46]. Specifically, we extracted ANN responses to a widely used fMRI localizer [42] which has also been used in prior studies of category-selectivity in ANNs [24–31, 36]. This dataset includes images of faces, bodies, scenes, objects, characters, and scrambled objects. To identify, say, face-selective units, we looked for units whose response to faces was significantly greater than to every other category at a stringent statistical threshold (*t >* 7, two-sided *p* = 7.17 × 10^−12^, see Methods). Next, we verified the selectivity of the identified units on an independent, publicly available stimulus set containing faces, bodies, scenes, and objects [24]. The procedure is shown schematically in **Fig. 2a**. The response profiles of the identified category-selective units (in the representative layer, see Methods) for stimulus categories in two representative ANN models (trained ResNet-50 ImageNet-1K and ViT-B/16 SigLIP2) are shown in **Fig. 2b, c**. The mean response to the preferred category (gray bar) was significantly higher than the response to all non-preferred categories (one-sided Mann-Whitney U tests; all *p* < 0.0001). We repeated the same analysis for 34 other pretrained ANN models (see Methods). These models span a range of architectures,training datasets, learning objectives, and model sizes (see **Supplementary Table 1** for the full list). Here too, the response to the preferred category was consistently and significantly higher than for other categories (one-sided Mann-Whitney U tests; all *p* < 0.05, **Fig. S1**) for all trained model architectures.

**Figure 2.**
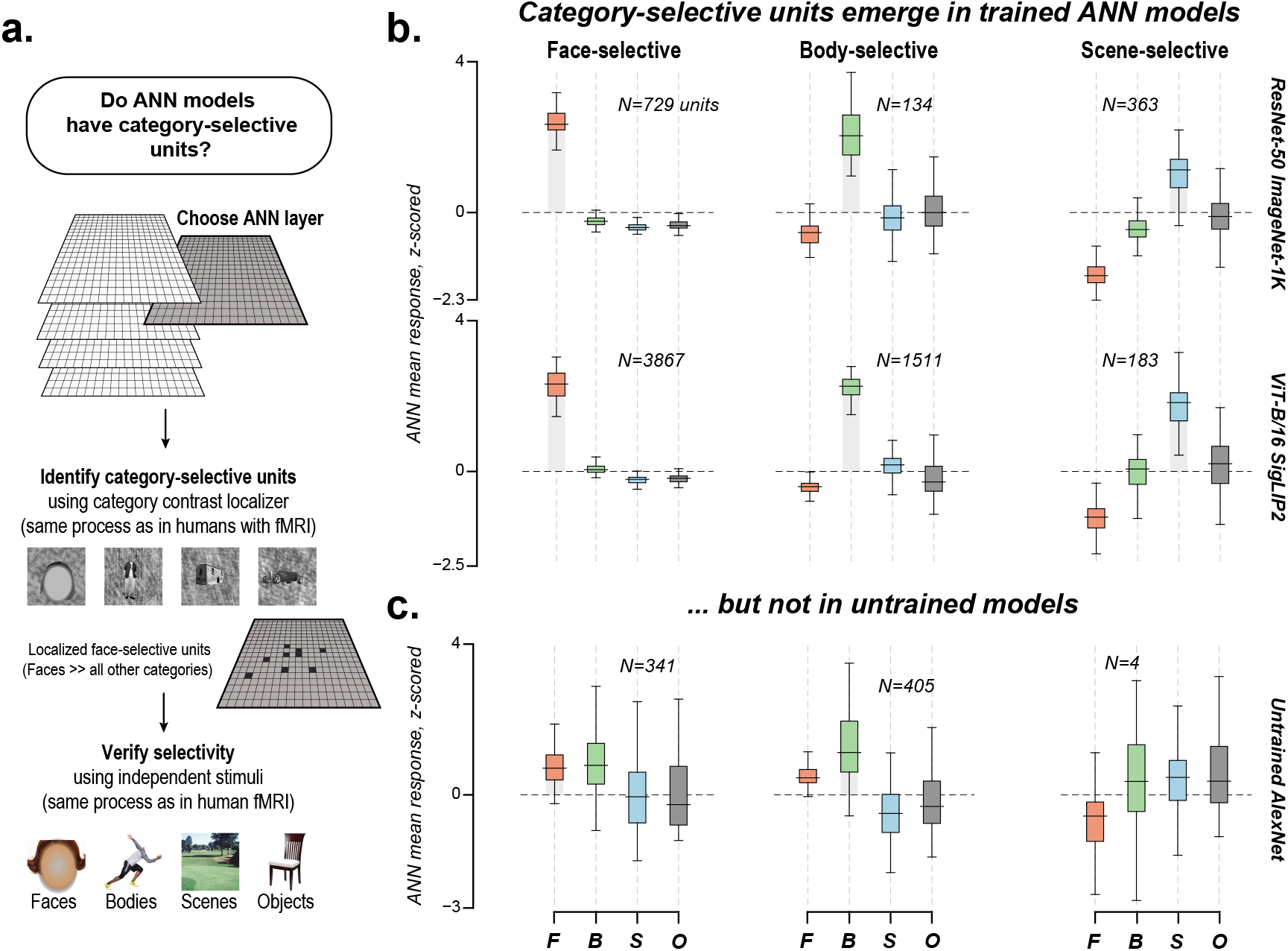
Trained, but not untrained, ANNs have reliable face-, body- and scene-selective units. **a** Schematic of the procedure used to identify category-selective units in a given ANN. For a layer (gray square), we apply a category-localizer [42] to identify candidate category-selective units, and validate their selectivity using an independent set of face, body, scene, and object images [24]. **b** Responses of localized category-selective units (from fROI-corresponding layer, *t >* 7; see Methods) to independent stimuli of faces (F), bodies (B), scenes (S), and objects (O). For all boxplots, the x-axis indicates the stimulus category, and the y-axis shows the unit-averaged z-scored responses. The text indicates the number of category-selective units localized. The first row is for a standard ANN model (ResNet-50 ImageNet-1K) and the second row is for the best-performing model (ViT-B/16 SigLIP2, see Methods). The columns show responses of face-, body-, and scene-selective units, respectively. **c** Same as **b** (except that t-threshold was lowered to *t >* 3, see Methods) but for an untrained AlexNet instance (seed= 5, see Methods).

Next, we asked if untrained (randomly initialized) ANNs also contain category-selective units. Note that our procedure of identifying category-selective ANN units is quite stringent and requires generalization to an independent stimulus set (as in human fMRI). We evaluated 20 untrained (randomly initialized) model instances (4 architectures × 5 seeds, see **Supplementary Table 2** for the full list) using the same procedure. None of the untrained models met the criterion for category selectivity across faces, bodies, and scenes at our originally defined statistical threshold (*t >* 7, **Fig. S2**). Note that some category-selective units could be detectable at lower selectivity thresholds (*t >* 3, two-sided *p* = 2.82 × 10^−3^). But the selectivity of these units did not consistently generalize to held-out stimuli (see **Fig. 2c** for an untrained AlexNet instance). These findings are in contrast with some prior claims of selectivity in untrained models [25] and show that model training is in fact essential for category selectivity in ANN units.

Taken together, these results show that category-selective units emerge reliably across a wide range of trained, but not untrained, ANN models. But the mere existence of category-selective ANN units that generalize to held-out images does not guarantee that the *form* of selectivity that arises matches what is observed in the human brain. We tested this next.

### 2.2 Category selectivity in ANNs is distinct from category selectivity in human brains: evidence from univariate measures

To meaningfully compare ANN selectivity to the brain, we needed to go beyond broad categories and characterize the full stimulus-level response tuning across a range of images and establish how reproducible those tuning patterns were across individuals (Q1, **Fig. 1b**). To address this question, we localized the category-selective voxels in the brain and measured the noise-corrected similarity of univariate response tuning across subjects in the FFA, EBA, and PPA using voxel-averaged responses to natural images from the Natural Scenes Dataset (NSD [41], see Methods). For each NSD subject (8 total), we correlated that subject’s voxel-averaged responses with those of each of the remaining seven subjects using a shared set of 515 images. The median of these seven correlations defined the subject-specific inter-subject ceiling. To account for measurement noise, correlations were normalized by the Spearman–Brown corrected within-subject split-half reliability (noise-corrected correlations can slightly exceed 1; see Methods). Responses in the canonical category-selective regions (FFA, EBA, and PPA) were strikingly consistent across individuals, with high noise-corrected human-human correlations (**Fig. 3b**; median *R* = 0.98 for FFA, 0.99 for EBA, and 1.01 for PPA; all individual brain–brain pair *R*s > 0.87, *p* < 0.0001). This cross-individual similarity of responses establishes a stringent benchmark for testing the response tuning found in category-selective ANN units (the NeuroAI Turing Test [47]).

**Figure 3.**
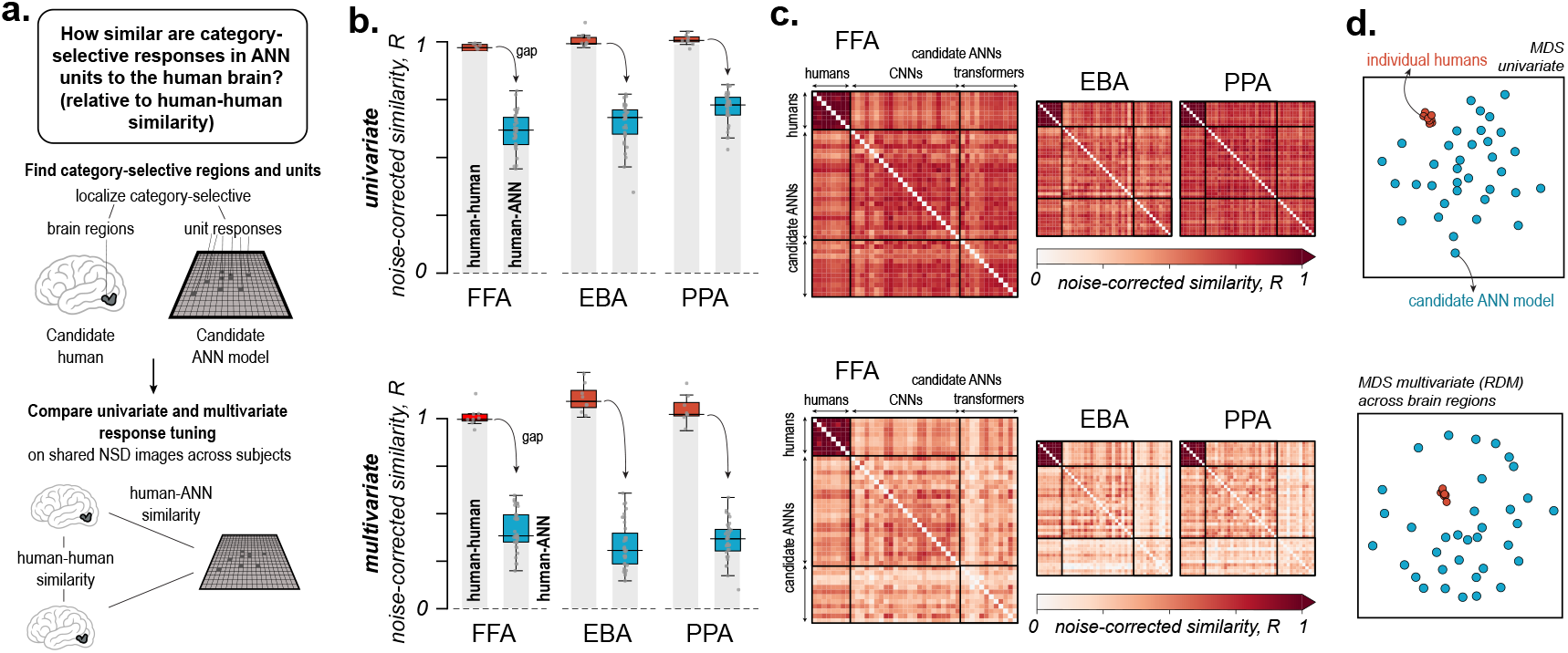
Category selectivity in ANN units is distinct from category selectivity in human brains by univariate and multivariate measures. **a.** Schematic of the analysis. Category-selective regions were identified in human fMRI data, and category-selective units were identified in ANN models using analogous localizer procedures. Responses to shared images from the Natural Scenes Dataset (NSD) were then compared between humans and ANNs using univariate and multivariate measures of similarity. **b.** Noise-corrected similarity between category-selective responses in humans and ANNs. For each region (FFA, EBA, PPA), boxplots compare human-human similarity (left) and human-ANN similarity (right). Each point denotes the median similarity for an individual subject (human-human) or ANN model (human-ANN). Top row, univariate comparisons. Bottom row, multivariate comparisons. Human-human similarity consistently exceeded human-ANN similarity across regions and metrics. **c.** Pairwise similarity matrices for category-selective response profiles. Subjects are arranged in the upper-left quadrant, ANN models in the lower-right quadrant, and human-ANN comparisons occupy the off-diagonal quadrants. Diagonal entries are masked. Darker colors indicate greater noise-corrected similarity. Top row, univariate comparisons. Bottom row, multivariate comparisons. **d.** Multidimensional scaling (MDS) visualization of pairwise similarities, averaged across face-, body-, and scene-selective regions. Each point represents a human subject (red) or ANN model (blue). Top row, univariate comparisons. Bottom row, multivariate comparisons. Human subjects cluster closely together, whereas ANN models occupy a distinct region of representational space.

#### Do ANN category-selective units exhibit human-like univariate response tuning?

We presented the same shared stimulus set viewed by the NSD subjects to each ANN model and measured the unit-averaged responses of the previously identified face-, body-, and scene-selective units (see Methods). Next, we correlated the ANN unit-averaged responses with the voxel-averaged responses measured from the human category-selective regions (noise-corrected as before, see Methods). Across all 35 pretrained ANN models and all three category-selective regions, human-ANN correlations were measurably and significantly weaker than human-human correlations (**Fig. 3b**; one-sided Wilcoxon signed-rank test on model-level differences, all *ps* < 0.00001). For example, the noise-corrected correlation between FFA responses and ANN face-selective responses was 0.62 (median across models), considerably lower than human-human FFA correlations (Δ*R* = 0.36, median across models). These correlations may seem high, so to provide additional context for the magnitude difference, we estimated the response similarity between the FFA and other category-selective regions like the OFA, EBA, and PPA. The gap between face-selective ANN units and the FFA was much larger than the gap between the FFA and OFA (Δ*R* = 0.06) and even the EBA (Δ*R* = 0.12) but not PPA (Δ*R* = 1.6) (**Fig. 3c, S3**). Note that the EBA is selective for an entirely different stimulus category (bodies). Thus, face-selective ANN units are less similar to the FFA than the tuning observed between several distinct human category-selective regions. The same pattern held for the EBA and body-selective ANN units (Δ*R* = 0.33) as well as the PPA and scene-selective ANN units (Δ*R* = 0.28), where the model–brain gaps exceeded those observed between different human category-selective regions (**Fig. 3b, S3**). Even the best ANN model we considered (ViT-B/16 SigLIP2) was significantly below the human–human ceiling (exact paired sign-flip permutation test across subjects, all one-sided *p*s < 0.05), with substantial gaps of Δ*R* = 0.32 for face selectivity, 0.25 for body selectivity, and 0.31 for scene selectivity. The full similarity pattern across all individual subjects and candidate ANN models sorted by convolutional and transformer architectures is shown in (**Fig. 3c**). To further visualize these differences across brain regions, we applied multidimensional scaling (MDS; 2 dimensions, see Methods) to the pairwise tuning curves across humans and ANNs (**Fig. 3d**). Human subjects formed a tight cluster, indicating high similarity with each other. ANN models (blue), on the other hand, were distinct from the human group (and to each other).

Together, these findings show that the voxel-averaged univariate stimulus-level response patterns in category-selective regions like the FFA, EBA, and PPA are remarkably consistent across human brains (in support of the second possibility for Q1, **Fig 1b**, left), whereas the unit-averaged category-selective responses observed in ANNs diverge significantly from those in humans (in support of the first possibility for Q2, **Fig 1b**).

### 2.3 Category selectivity in ANNs is distinct from category selectivity in human brains: evidence from multivariate measures

It remains possible that the representations in category-selective ANN units more closely resemble human category-selective representations when considering the multivariate pattern of responses (beyond the voxel/unit-averaged univariate tuning) [48–53]. To directly test this possibility, we compared the multivariate structure between ANNs and brain responses using representational dissimilarity matrices (RDMs) (see Methods). We constructed RDMs based on Euclidean distances between pairs of images across voxels/units (appropriate for categorical stimuli, see [54]). As in the univariate analyses, we accounted for measurement noise by normalizing the pairwise RDM correlations with each subject’s split-half RDM reliability across repeated stimulus presentations. We first asked how consistent the multivariate structure of responses was across human subjects. As for the univariate analyses, we found that RDMs from the FFA, EBA, and PPA were highly similar across participants (**Fig. 3b**; median *R* = 0.99 for FFA, 1.09 for EBA, and 1.02 for PPA; all individual human-human pair *Rs*>0.81, *p* < 0.0001). These results show that even the fine-grained multivariate response patterns in category-selective cortex are remarkably stable across individuals (further support possibility 2 for Q1).

#### Do category-selective units in ANNs exhibit multivariate response structure similar to that observed in human category-selective brain regions?

We compared the representational structure of ANN face-, body-, and scene-selective units to that of the FFA, EBA, and PPA in humans. As before, we used the same images to measure noise-corrected correlations between RDMs observed in ANN and in brains. Across all models and category-selective regions, human-ANN correlations were significantly lower than the ceiling established by human-human correlations (**Fig. 3b**, one-sided Wilcoxon signed-rank test on model-level differences, all *ps* < 0.00001). For the FFA, the median gap between response tuning similarity with ANN models (Δ*R* = 0.64) was again higher than the similarity with OFA (Δ*R* = 0.05) and EBA (Δ*R* = 0.19) but not PPA (Δ*R* =0.56) (**Fig. 3c, S4**). Similarly, the median gap (Δ*R*) between human-human and human-ANN correlations was 0.78 for EBA and 0.68 for PPA. Strikingly, these gaps were larger than those for univariate tests (**Fig. 3b**). Even for the best-performing model (ViT-B/16 SigLIP2), the human-ANN correlation was significantly below human-human ceiling (exact paired sign-flip permutation test across subjects, all one-sided *ps* < 0.05) with a gap (Δ*R*) of 0.43 for face-selectivity, 0.62 for body-selectivity, and 0.75 for scene-selectivity. We also visualized the multivariate response across models and subjects’ similarity with MDS (**Fig. 3d**). As expected,human subjects clustered closely together, showing very similar response patterns, whereas ANN models were more spread out and different from one another.

To summarize, we find that category-selective responses in the human brain are remarkably consistent across individuals by both univariate and multivariate measures. In contrast, category-selective units in ANNs differ both in their overall univariate response tuning and in the structure of their multivariate representations. Even when ANN units have the same category preference as human category-selective regions, their response patterns are measurably different. This suggests that even though category-selective units indeed exist in ANNs, their patterns of responses are highly variable across models and not brain-like.

### 2.4 Other factors do not explain differences between category-selectivity in ANNs and human brains

We identified three key experimenter-chosen parameters that could impact the comparison between category-selectivity in ANNs and human brains: 1) the selectivity threshold used to identify category-selective units and voxels, 2) the functional localizer used to identify units within ANNs, and 3) the choice of brain category-selective fROI used as the comparison target for ANN category-selective units. Next, we systematically assessed the impact of each factor on human-ANN similarity and showed that the overall pattern of results remains unchanged.

#### Category-selectivity threshold

Applying a more stringent selectivity threshold could, in principle, yield ANN units whose responses more closely match human category-selective regions. To test this possibility, we systematically varied the selectivity thresholds used to identify category-selective ANN units and human voxels. As expected, higher thresholds produced fewer but more selective units and voxels (**Fig. 4e**). However, across all threshold combinations and all three regions of interest, human-ANN correlations remained significantly below the human-human ceiling for both univariate tuning (**Fig. 4f**; one-sided Wilcoxon signed-rank test on model-level differences, all *ps* < 0.00001) and multivariate representational structure (**Fig. 4f**; one-sided Wilcoxon signed-rank test on model-level differences, all *ps* < 0.00001). Even the best-performing model (ViT-B/16 SigLIP2) did not approach the human-human ceiling across any combination of selectivity thresholds (**Fig. S5**; exact paired sign-flip permutation test across subjects, all one-sided *ps* < 0.05; lowest achieved median multivariate ΔR = 0.41, 0.56, and 0.69 for face-, body-, and scene-selectivity respectively). These results indicate that the observed ANN–brain differences cannot be explained by the selectivity threshold used to define category-selective units or voxels, and that the representational gap persists across a wide range of selectivity strengths.

**Figure 4.**
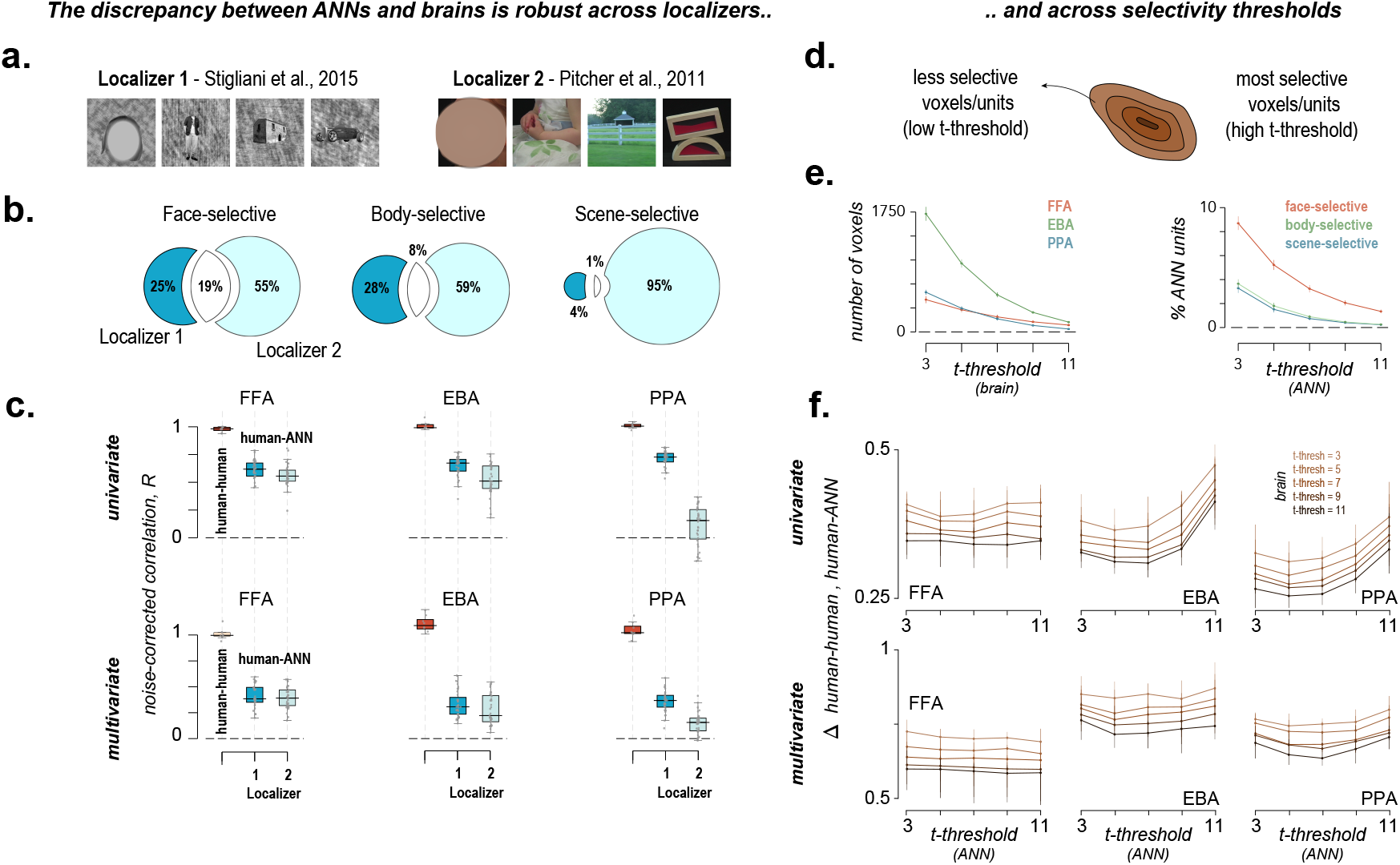
The discrepancy between selectivity in ANN units and human brains is robust across localizers and selectivity thresholds. **a.** Example images from two different functional localizers used to identify category-selective units [42, 43]. **b.** Overlap between category-selective ANN units identified by the two localizers. Percentages indicate the proportion of units unique to each localizer or shared between localizers (median across models). **c.** Human-human and human-ANN similarity for category-selective responses identified using each localizer. Human-human similarity consistently exceeded human-ANN similarity across localizers, regions, and both univariate (top) and multivariate (bottom) analyses. Each point represents one subject (human-human) or one ANN model (human-ANN) pair. **d.** Schematic illustrating the relationship between selectivity strength and the t-threshold used to define category-selective voxels and units. **e.** Effect of t-threshold on the number of category-selective voxels in the brain (left) and the proportion of category-selective units in ANNs (right). Increasing threshold (x-axis) identifies progressively more selective and fewer voxels and units (y-axis). Colors indicate each kind of selectivity. **f.** Gap between human-human and human-ANN similarity (y-axis) as a function of voxel and unit selectivity thresholds (x-axis). The discrepancy between ANN and brain category selectivity remained robust across a broad range of threshold choices for both univariate (top) and multivariate (bottom) analyses.

#### Functional localizer

It’s also possible that our initial localizer simply failed to identify the most brain-like category-selective units in ANNs. To rule out this possibility, we compared units identified using an alternative, widely adopted fMRI localizer [43] to those selected with the original localizer used in human studies [42]. Strikingly, the two unit subsets exhibited low overlap across all categories (**Fig. 4b**; median overlap (IoU) = 18.8% for faces, 9.3% for bodies, and only 1.5% for scenes; see methods). This overlap is significantly lower than the within-localizer split-half overlap seen in the brain (**Fig. S6**; one-sided Wilcoxon signed-rank test, all *ps* < 0.0001; see Methods). This result contrasts with the human brain, where category-selective regions can be reliably identified across different localizers [46]. In ANNs, by contrast, category selectivity was highly sensitive to the particular stimulus set used to localize them (which must be investigated more systematically in future work). But does this alternative localizer yield units that better match the brain? We found that it does not. Across all ANNs and category-selective regions, human-ANN correlations remained significantly below the human-human ceiling for both univariate and multivariate comparisons in FFA, EBA, and PPA (**Fig. 4c**; one-sided Wilcoxon signed-rank test on model-level differences, all *ps* < 0.00001). This raised another critical concern. Because different localizers identify entirely different subsets of category-selective units (a breakdown in its own right), it is possible that ANN models do contain other unit subsets that are brain-like. We tested this possibility directly by dropping the category-selectivity constraint and instead selecting, for each voxel, the ANN unit whose responses were most strongly correlated with that voxel [55] (see Methods). Even under this more permissive procedure, human-ANN correlations were significantly below the human-human for both univariate and multivariate comparisons in FFA, EBA, and PPA (**Fig. S7**; one-sided Wilcoxon signed-rank test on model-level differences, all *ps* < 0.00001). The gap remained substantial across univariate (median Δ*R* = 0.22, 0.26, and 0.22 for face-, body-, and scene-selectivity, respectively) and multivariate comparisons (median Δ*R* = 0.5, 0.68, and 0.61 for face-, body-, and scene-selectivity, respectively). Thus, even the most brain-aligned unit subsets identifiable within current ANN models failed to reproduce the stable stimulus-level category-selective response patterns observed in human brains.

#### Choice of category-selective fROIs

Another possibility is that category-selective ANN units may align more closely with *other* category-selective regions beyond than FFA, EBA, or PPA. To test this possibility, we compared face-, body-, and scene-selective ANN units with a broader set of 5 other category-selective fROIs (OFA, FBA, OPA, VWFA, and OVWFA, see Methods). Across all fROIs, the human-ANN correlations were significantly lower than human-human ceilings across both univariate and multivariate comparisons (**Fig. S8**; one-sided Wilcoxon signed-rank test on model-level differences, all *ps* < 0.00001). Thus, category-selective ANN units are not brain-aligned even when comparisons extend beyond the canonical category-selective regions.

Taken together, these results show that the difference between ANN and human category selectivity is robust across analysis choices and extends beyond FFA, EBA, and PPA.

### 2.5 Category-selective ANN units are neither necessary nor sufficient for predicting brain responses

So far, our results show that category-selective ANN units do not display the stable univariate and multivariate response profiles observed in human category-selective brain regions. At the same time, we know from prior studies that ANN features can be linearly combined to predict responses in human visual cortex [56–59], including category-selective regions such as the FFA, EBA, and PPA [24, 45, 60]. One possibility is that category-selective units, even though not similar to category-selective voxels directly, are critical to the prediction accuracy of encoding models [61]. We tested this idea directly by asking: *Are category-selective units in ANNs necessary and/or sufficient for predicting category-selective responses in the brain?*

To test necessity, we performed a lesioning manipulation by removing all the previously identified category-selective ANN units (sections 2.1-2.3) and then building voxel-wise encoding models to predict responses in each target region (FFA, EBA, and PPA). We reasoned that if category-selective units were necessary for prediction, removing them should reduce model prediction scores relative to encoding models trained on all units in the layer. To test sufficiency, we trained models using *only* the category-selective units as input features. If these units were sufficient, models trained only on category-selective units should match the predictive accuracy of models trained on all units in the layer. Note that the encoding models were trained on an independent set of 485 images (shared across 4 subjects) and prediction accuracy was evaluated on the 515 held-out images used in all previous analyses (see Methods).

We found that category-selective units were neither necessary nor sufficient for the predictive accuracy of the encoding models (**Fig. 5**). Lesioning category-selective units had minimal impact on predictive accuracy with lesioned models performing nearly identically to full models across all three regions (**Fig. 5b, c**). Lesioning selective units reduced performance for univariate comparisons by 0.22% [95% CI: 0.10%, 0.26%] with respect to the within-subject ceiling, and for multivariate comparisons by 0.47% [95% CI: 0.28%, 0.79%] (median across models, fROIs, and subjects). Moreover, models trained *only* on category-selective units performed substantially worse than models trained on all units with a performance drop of 18.37% [95% CI: 15.45%,24.14%] for univariate comparisons, and 30.58% [95% CI: 28.49%, 37.98%] for multivariate comparisons (**Fig. 5b, c**, median across models, fROIs, and subjects), showing that these units are not sufficient on their own. Our findings were robust and hold across different selectivity thresholds (**Fig. S9**) and under a constrained sparse-positive mapping between ANN units and the brain [24, 62] (**Fig. S10**).

**Figure 5.**
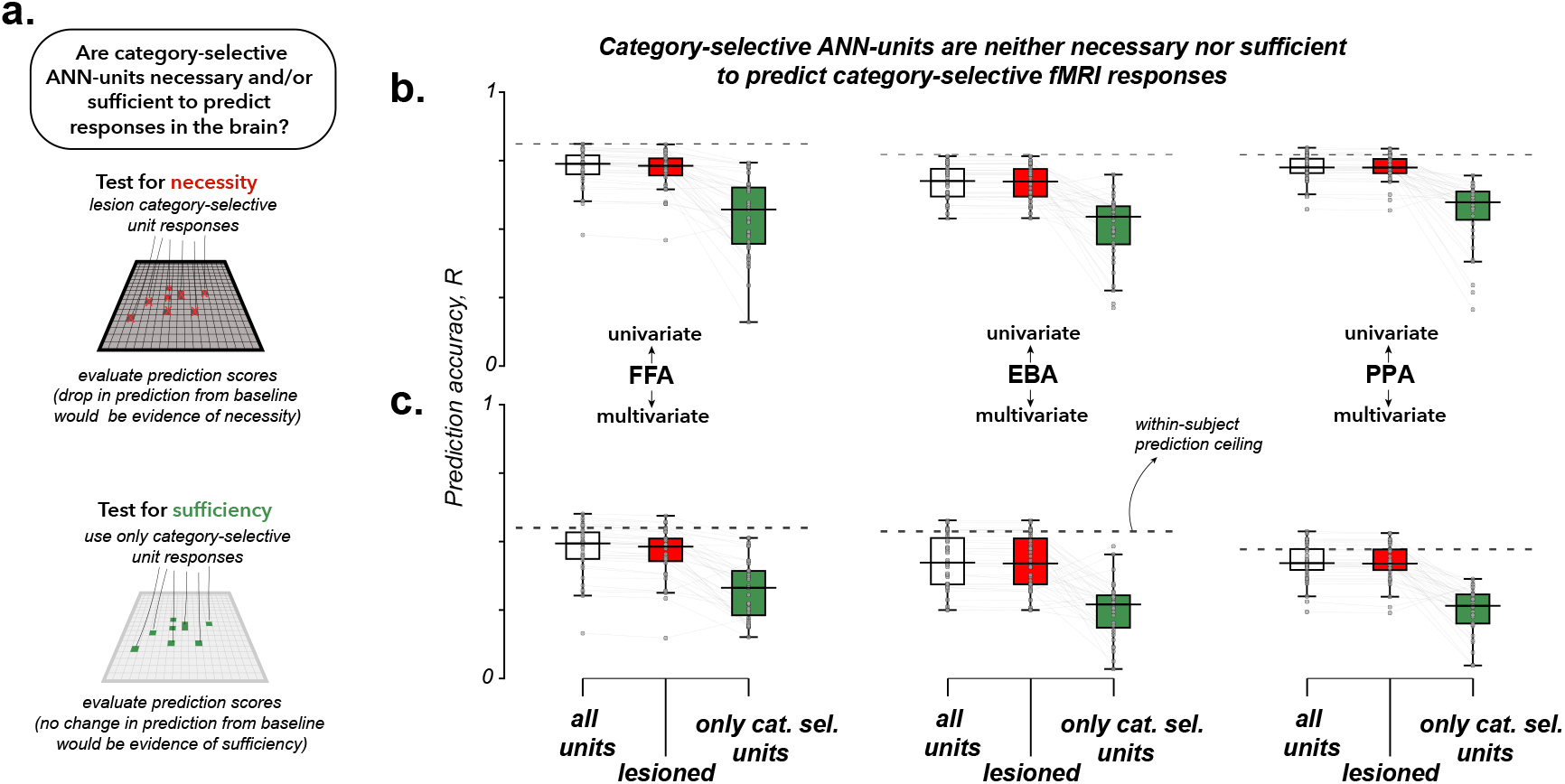
Voxel-wise encoding model predictivity from ANN units to category-selective regions is minimally impaired when ANN category-selective units are lesioned. **a** Schematic describing the question and tests to investigate whether ANN category-selective units drive the predictive accuracy of voxel-wise encoding models of category-selective regions. **b** Correlation between observed and predicted responses from voxel-wise encoding models (using ridge mapping) on held-out NSD images. For each boxplot, the x-axis indicates the unit set considered, and the y-axis indicates the correlation. The first box depicts the correlation of the encoding model using all the units within the ANN layer (each point indicating one ANN model’s median correlation across the subjects); similarly, the second and third boxes correspond to encoding models when the category-selective units are lesioned, and when only category-selective units are considered. The black dashed line shows the median within-subject ceiling, i.e., spearman-brown corrected split-half correlation between the subject’s responses during the three repetitions of the same stimuli. The two rows indicate univariate and multivariate comparisons, respectively. The three columns indicate the category-selective region, i.e., FFA, EBA, and PPA, respectively.

Together, these findings provide an independent line of evidence that category-selective ANN units do not instantiate category selectivity in the same way as the human cortex. Category-selective units in ANN models were neither necessary nor sufficient for predicting responses in human category-selective brain regions. Information necessary for brain predictivity of category-selective regions appears to be distributed broadly across ANN representations rather than concentrated within ANN units deemed category-selective.

### 2.6 Interpreting the differences between category-selectivity in ANNs and fMRI voxels

Our analyses so far show a quantitative gap between category selectivity in ANN units and brain regions. But aggregate statistical metrics do not tell us anything human-interpretable about *which* aspects of the stimulus structure are responsible for these differences. Both ANNs and brains represent information across many interacting stimulus dimensions [58, 63–69], and the differences likely arise from a combination of factors. Here, we begin to unpack this complexity by identifying some interpretable stimulus conditions patterns that elicit systematic response differences between category-selective ANN units and brain regions.

This analysis is particularly prone to circularity [70], and therefore we developed a strict data-partitioned hypothesis-generation and testing framework (summarized in **Fig. 6a**). We developed our hypotheses by examining the responses from a single standard model (ResNet-50 ImageNet-1K) and a single participant (S1 from NSD). Using this model-subject pair, we identified images that produced especially large response differences between ANN unit-averaged responses and brain voxel-averaged responses. We used these outlier images to infer the stimulus-level factors that may differentiate ANN unit and brain category-selective responses. Crucially, this exploratory phase was completely separated from the hypothesis testing. The hypotheses were evaluated on entirely independent fMRI data from held-out subjects and from images drawn from the 9000 held-out subject-specific images from the NSD. All the hypotheses were tested across the top 10 ANN models whose representations most closely matched human category-selective regions (see **Supplementary Table 1** for the list of models). This strict separation between hypothesis development and validation allowed us to assess whether the observed effects generalized to (1) held-out subjects (N=7 participants), (2) held-out stimuli (N=350 images), and (3) held-out models (N=10 models). As expected, not all our initial hypotheses held up in independent tests, and we report all outcomes below.

**Figure 6.**
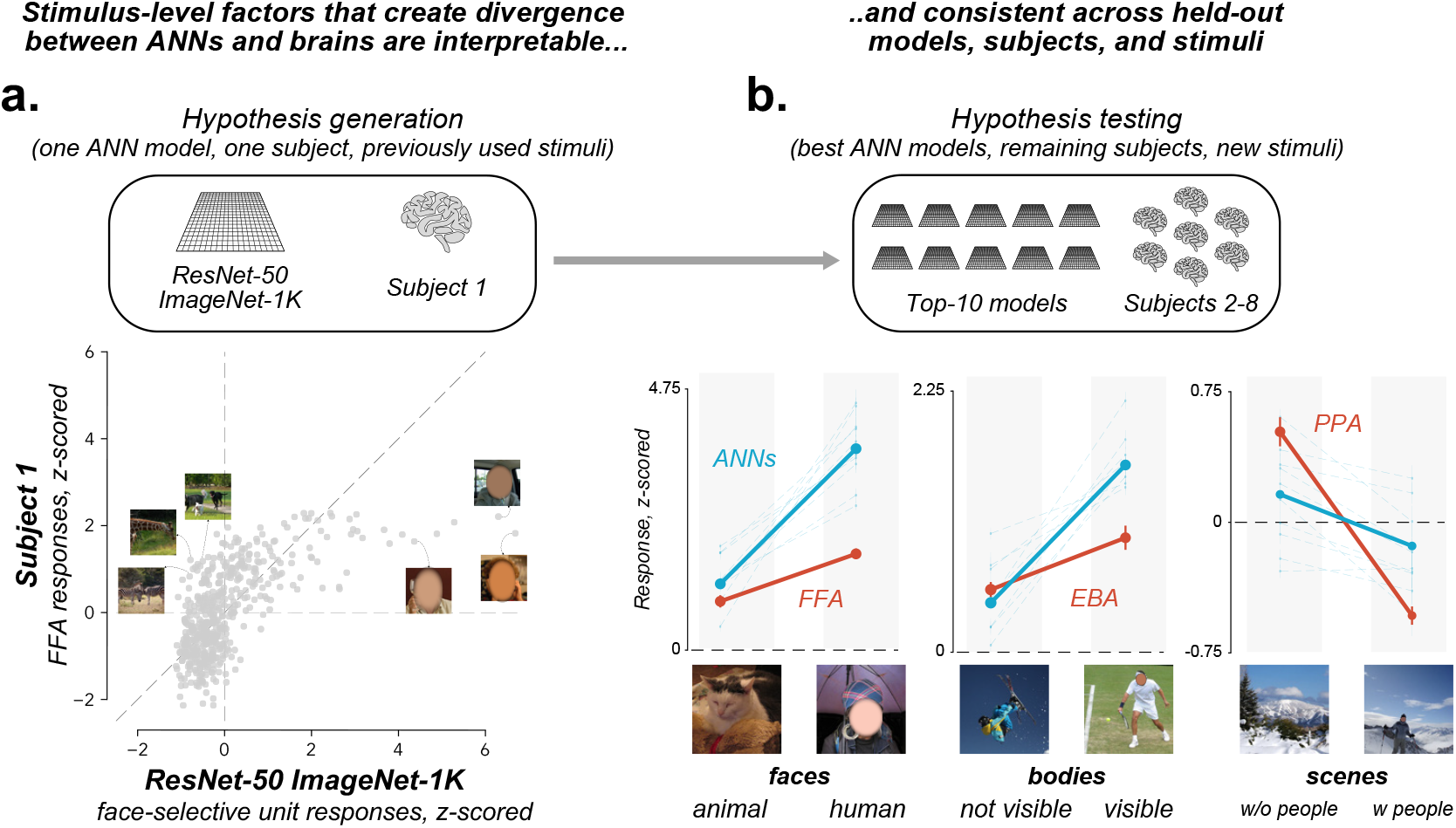
Stimulus-driven divergences between ANN and human category-selective responses generalize across held-out models, subjects, and stimuli. **a** Schematic of the hypothesis-generation framework. Hypotheses were generated using responses to the previously used evaluation subset (shared 515 NSD stimuli), from a single reference subject (S1) and a single standard model (ResNet-50 ImageNet-1K). The lower panel visualizes the identification of divergent stimuli for face selectivity. Divergent (outlier) stimuli were identified based on discrepancies between unit-averaged face-selective ANN responses (x-axis) and voxel-averaged FFA responses (y-axis), both z-scored across stimuli. NSD images are from the COCO image dataset/Flickr [74]. **b** The top panel shows a schematic of the hypothesis-validation framework. The candidate stimulus-level factors were evaluated on independent, subject-specific held-out images, held-out subjects, and held-out top-10 most brain-aligned models according to univariate tests (see Methods). The middle panel shows example NSD images from the stimulus groups that produced systematic divergence between ANN and brain responses across three domains of category selectivity: face-selective (animal faces vs. human faces), body-selective (bodies with obscured limbs vs. visible limbs), and scene-selective (scenes without humans vs. scenes with humans). The bottom panel shows a descriptive visualization of ANN–brain divergence for each stimulus group. The x-axis indicates stimulus group, and the y-axis shows unit-averaged (ANN) or voxel-averaged (brain) z-scored responses (see Methods). Three columns show divergences for face-, body-, and scene-selectivity, respectively. For each plot, the thick red lines denote the mean response across subjects (points indicate mean *±* SEM), thick blue lines denote the median response across the models, and faint blue lines show individual model responses. Across all domains, ANNs exhibit robust differences between stimulus groups relative to human category-selective regions. NSD images are from the COCO image dataset/Flickr [74].

#### ANN face-selective units overemphasize the distinction between human and animal faces

The clearest and most interpretable differences between ANN and brain responses emerged in the domain of face selectivity. In the exploratory phase (using the single reference model and subject), we found that face-selective ANN units showed an exaggerated difference between human and animal faces compared to the FFA (**Fig. 6b**). While the FFA did produce a stronger response to human than animal faces, the difference was substantially and disproportionately larger in ResNet-50 face-selective units. This led to a concrete hypothesis: ANN face-selective units systematically overemphasize the distinction between human and animal faces compared to the human FFA (**Fig. 6b**). We found strong support for our hypothesis on held-out subjects, stimuli, and models (see above). For each subject, we curated independent images containing either human faces or animal faces (350 stimuli total, 25 stimuli per group per subject). As reported in other studies [71, 72], we found that the FFA responses were relatively higher to images containing human faces than animal faces (**Fig. 6c**; Linear mixed-effects model, simple effect of group for FFA: all *z* > 4.8, one-sided *p* < 0.00001; see Methods for full model specification). While ANNs also exhibited higher responses to human faces than animal faces (simple effect of group for ANNs: all *z* > 14.5, one-sided *p* < 0.00001), the overall magnitude of this difference was significantly larger across all models (**Fig. 6c**; 10 out of 10 models, group x system interaction: all *z* > 4.5, one-sided *p* < 0.00001). As a negative control, we replicated the same analyses using randomly sampled images (N=100 bootstraps; for each bootstrap, 350 stimuli total, 25 stimuli per group per subject to match the experiment) and found no differences (**Fig. S13**). Together, these results indicate that face-selective ANN units exaggerate the difference between human and animal faces compared to the human FFA, and may broadly overemphasize visual cues related to human faces.

#### ANN body-selective units are highly sensitive to limb visibility

Based on our previous result with faces, it was natural to ask if there was a similar distinction between ANN units and the EBA for human and animal bodies. However, we found limited evidence for systematic differences under our stringent hypothesis-testing framework (**Fig. S12**). Instead, somewhat surprisingly, a different, more consistent pattern emerged with respect to limb visibility (**Fig. 6b**). In our exploratory analyses, ANN body-selective units appeared particularly sensitive to whether human limbs were visibly exposed versus obscured by heavy clothing. This led to a new hypothesis: ANN body-selective units rely more heavily on low-level visual cues (like skin visibility/color) than the human EBA. As before, we curated independent images from each subject’s unique images containing human bodies with prominent limbs versus not (350 total stimuli, 25 per group per subject). We found that EBA responded slightly more to bodies with visible limbs than fully-covered bodies (**Fig. 6c**; Linear mixed-effects model, simple effect of group for EBA: all *z* > 5.4, one-sided *p* < 0.00001), consistent with prior work [73]. But as hypothesized, the reliance on limb-related low-level cues was considerably higher in ANN units (**Fig. 6c**; 9 out of 10 models, group x system interaction: all *z* > 2.3, one-sided *p* < 0.05). As a negative control, randomly sampled images exhibited no divergence (**Fig. S13**). Together, these results suggest that ANN body-selective units place disproportionate weight on surface-level visual cues such as exposed skin, whereas the EBA exhibits a more graded and less cue-dependent representation of bodies.

#### Human scene-selective regions, and not ANN scene-selective units, are strongly suppressed by humans in scenes

In exploratory analyses, we observed that human scene-selective responses in the PPA were strongly suppressed by the presence of people within scenes (**Fig. 6b**). This led to a clear hypothesis: scene-selective units in ANNs show weaker suppression to humans in scenes compared to the PPA. Note that we also explored additional hypotheses (indoor versus outdoor scenes, scenes with lower versus higher navigational affordances), but these hypotheses did not produce consistent or reliable differences between ANN scene-selective units and PPA (**Fig. S12**). To test our hypothesis, we curated images of scenes with and without humans in them (350 total stimuli, 25 per group per subject). The PPA responded significantly less to scenes containing humans than to scenes without humans (**Fig. 6c**; Linear mixed-effects model, simple effect of group for PPA: all *z* > 11.5, one-sided *p* < 0.00001). ANN scene-selective units showed the same qualitative pattern, but as hypothesized, the magnitude of the suppression was substantially smaller in ANNs. Across all 10 tested models, there was a significant group × system interaction (all *z* > 3.1, one-sided *p* < 0.005), indicating that the attenuation of responses by human presence in scenes was reliably weaker in ANNs than in the PPA (**Fig. 6c**). Additionally,randomly sampled images showed no difference (**Fig. S13**). Overall, while both scene-selective ANN units and the PPA are modulated by humans within scenes, the PPA exhibits a much stronger suppression effect. In contrast, ANN scene-selective units appear less sensitive to this contextual modulation.

To summarize, these results expose consistent and interpretable differences between category selectivity in ANNs and brains. Across faces, bodies, and scenes, category-selective ANN units appear strongly influenced by the specific visual features present in the localizer stimuli. In contrast, category-selective responses in the brain generalized more consistently across stimuli and individuals. These findings suggest that category selectivity in current ANN models is comparatively fragile, relying more heavily on the particular cues used to define a category, whereas category selectivity in the brain reflects a more stable and shared representational organization.

## 3 Discussion

In this work, we draw the distinction between the *existence* of category selectivity in modern ANNs and the *form* of category selectivity they exhibit compared to the human brain. We first show that the stimulus-level response tuning in category-selective regions is highly consistent across individuals (*P*_2_ for Q1, **Fig. 1**), and can be used as a representational benchmark for ANNs. We then applied the same localization procedure used in human fMRI to ANNs and found that face, body, and scene-selective units emerge reliably in trained, but not in untrained models (**Fig. 2**). Next, we compared the detailed stimulus-level response patterns and found that category-selective ANN unit responses differed substantially (both in univariate tuning and multivariate response structure, **Fig. 3**; *P*_1_ for Q2, **Fig. 1**) relative to the human FFA, PPA, and EBA. Our results were robust to analysis choices like the functional localizer used, the selectivity threshold, and the specific regions examined (**Fig. 4**). Although ANN features could be linearly combined to predict voxel responses in category-selective cortex (using encoding models), lesioning experiments further confirmed that the category-selective ANN units themselves were neither necessary nor sufficient for this predictivity (**Fig. 5**). Finally, we used stimulus-level tests to expose some interpretable differences between category-selective ANN units and brains (**Fig. 6**). These findings suggest that the representations in category-selective ANN units are driven by superficial, dataset-dependent cues and differ in important ways from the consistent and structured representations observed in human category-selective brain regions.

The first key finding from our study concerns the brain itself. Category-selective regions are usually defined with coarse functional localizers that use only a small number of images and stimulus categories (typically two to five). This process raises a natural concern that the localizers might simply be grouping random voxels that pass a category contrast without otherwise a shared representational structure [75–77]. Our results suggest otherwise: category-selective regions of the brain show highly similar stimulus-level response patterns across individuals. This representational stability across individuals has several key implications. From a neuroscience perspective, it means that the fROI-based method can be used to find stable and meaningful differences between different regions (e.g., FFA vs. OFA [43, 78]), developmental stages (e.g., children vs. adults [22, 79–82]), and species (e.g., human FFA vs. ML/AL patch in macaque inferotemporal cortex [83, 84]). From a modeling perspective, this stability enables a direct comparison of selectivity in ANNs and brains. In fact, unlike the stable category selectivity observed in the human brain [46], different localizers identified very different subsets of selective units in ANNs. This suggests that selectivity in current ANN models is considerably more fragile and relies overly on the specific features present in the images used to probe the model. This observation leads to two practical recommendations for future NeuroAI research. First, studies comparing ANN units to brain regions should localize those that mirror those used in human neuroscience. Second, they should demonstrate that conclusions remain stable across different localizer choices.

The second (and central) finding of our study is that category selectivity in ANN units cannot (yet) be directly equated with the category selectivity observed with human fMRI. As we detailed in our Introduction, a category-contrast is an underconstrained test because widely different response patterns can easily satisfy it. From a model-to-brain alignment perspective, it calls for caution because the overriding category-level variance can make two systems look more similar than they really are, unless one uses more demanding analyses like comparing the noise-corrected image-grained response profiles. Therefore, finding category-selective units in ANNs should only be considered the first step. When we compared the full image-level response structure, many differences became clear. ANN units did not approach the stable level of similarity observed across human brains, either in univariate tuning or in multivariate representational structure. In many cases, ANN units were less similar to a target brain region (e.g., FFA) than other, functionally distinct category-selective regions were (e.g., EBA). Category-selective units were also highly variable across ANN models, suggesting that different models in fact converged on different solutions rather than a common brain-like form of selectivity. These mismatches extended beyond the FFA, EBA, and PPA to the broader set of category-selective regions and remained robust across localizers and selectivity thresholds. Consistent with this observation, category-selective ANN units were neither necessary nor sufficient for predicting responses in human category-selective cortex: removing these units had little effect on prediction scores, whereas restricting models to these units substantially reduced model predictivity. Taken together, these results show that ANNs can satisfy the minimal definition of category selectivity without reproducing the stable and shared form of selectivity observed in the human brain with fMRI.

Our findings open several important questions. First, what evolutionary and developmental pressures give rise to brain-like category representations? We find that ANNs trained on a wide range of tasks do not yet converge on the form of category selectivity observed in the human brain. As other studies have shown, ANNs are brain-aligned at a representational level, often more so than previous generations of models [60, 63, 85–87]. Future work will need to ask what additional constraints are required for brain-like organization to emerge. Prominent ideas suggest developmental constraints [88–91], ecological visual experience [92–95], wiring and topographic pressures [27–32], recurrent processing [96, 97], learning rules, and objectives that better capture the problems the visual system evolved to solve. Second, what determines the stability of category-selective representations? As we show, category-selective responses are highly stable across people and depend on more robust features (e.g., human vs. animal faces in FFA) [98]. As we also show, category-selective responses are much more fleeting in ANN units, often overspecialized to the features in the localizer images used to identify them. This is concerning because it shows that ANNs do not share biological invariances. Our study also gives practical guidance for how to probe the stability of these models in interpretable ways. Even simple contrasts, like human versus animal faces, scenes with versus without people, or bodies with visible versus covered limbs, are enough to reveal clear differences between voxels and ANN and can be used as diagnostic tests. Future studies that claim to find a better match between ANN units and brains should directly use these tests. Finally, our work forces us to think about the appropriate level of analysis for comparing brains and ANNs. Our results suggest that, at least in current ANN models, individual units may not be aligned to brains. Combinations of units in the meantime might be better suited for using ANN models as tools for understanding the brain [24, 45, 60, 99]. This pattern fits with prior work showing that unit-level metrics (such as RSA) often yield a weaker match to the brain than encoding models that combine features across several units [100, 101]. Future work should continue to study both levels and develop better, sharper metrics [102–105]. Population-level analyses may currently provide the strongest signal of alignment, but it will also be important to test more directly whether individual artificial units can match the response tuning of single voxels, and ultimately single neurons.

To summarize, we show that category selectivity is a much stronger test of ANN models than it initially appears. Even though current ANN models contain units that can be deemed category-selective, the form of category selectivity they exhibit remains consistently and measurably different from that observed in the human voxels.

## 4 Methods

### 4.1 fMRI data and localization details

We analyzed data from the 1.8 mm^3^ native surface preparation of the Natural Scenes Dataset (NSD; [41]), using version 3 single-trial beta estimates (betas_fithrf_GLMdenoise_RR). To reduce session-to-session variability, voxel responses were z-scored within each session and then averaged across the three repetitions of each stimulus. We focused on the 1000 images shared across subjects. Of these, 515 images were viewed three times by all 8 subjects and served as the main evaluation set. The remaining 485 images were viewed three times by a subset of 4 subjects (1, 2, 5, and 7) and were used as a training set for selecting representative ANN layers, selecting best-performing models, and fitting voxel-wise encoding models.

Our primary functional regions of interest (fROIs) included the fusiform face area (FFA; combining FFA-1 and FFA-2), extrastriate body area (EBA), and parahippocampal place area (PPA). Voxels were selected bilaterally based on category-selective responses from an independent functional localizer [42], using a contrast-specific t-value threshold. Unless otherwise noted, we used a threshold of *t >* 7.

### 4.2 Modeling details

#### 4.2.1 Artificial Neural Network models

We analyzed 35 task-optimized artificial neural network (ANN) models spanning a diverse set of architectures, training datasets, learning objectives, and model sizes. Models were sourced from several widely used repositories, including the Torchvision (PyTorch) model zoo [106], PyTorch Image Models (timm; [107]), OpenCLIP [108], OpenAI CLIP [109], the Open-IPCL project [110], and Huggingface Transformers [111]. A complete list of models is provided in **Supplementary Table 1**.

To assess the role of training, we additionally analyzed 20 randomly initialized (untrained) models. These consisted of four standard architectures (AlexNet, ResNet-18, ResNet-50, and ViT-B/32), each instantiated with five different random seeds (1–5). Model weights were initialized using the default PyTorch initialization procedures. The full list of untrained models is provided in **Supplementary Table 2**.

To extract ANN unit responses, we presented stimuli to each model and recorded activations from a given intermediate layer. Each image was first zero-padded to a square along the shorter dimension to preserve aspect ratio, then processed using the model’s native inference transforms.

### 4.2.2 Identifying category-selective units within an ANN layer

To make the comparison with the brain as fair as possible, our goal was to use the same localization logic in ANNs that is used to identify category-selective regions in human fMRI. We therefore identified category-selective units by presenting a standard functional localizer. Our main analysis used the vpnl-fLoc localizer [42], which contains grayscale images with low-level properties matched across categories and includes 288 images each of faces, scenes, bodies, objects, and characters, along with 144 scrambled images. Note that this is the same localizer used in the Natural Scenes Dataset fMRI data used in our study [41].

We identified category-selective units by testing whether each unit responded more strongly to a preferred category than to every other category separately. For example, a face-selective unit had to respond more strongly to faces than to scenes, bodies, objects, characters, and scrambled stimuli. For each comparison, we computed a two-sample t-test, and units that exceeded the threshold in all comparisons were classified as category-selective. We repeated this procedure for face-, scene-, and body-selective units. Our main analysis used a threshold of *t >* 7 (two-sided *p* = 7.17 × 10^−12^, *df* = 574) for pretrained models, which was matched with the fMRI data. We report the corresponding p-values to indicate the stringency of the threshold; the threshold was used as a selection criterion to define category-selective unit populations rather than to perform statistical inference across individual units. Note that all analyses were conducted at the exemplar (single-stimulus) level, such that unit responses were evaluated on individual images rather than block-averaged category responses. This differs from standard fMRI localizer analyses, which typically rely on category-level contrasts (e.g., faces vs. non-faces in block designs); here, selectivity was defined at the level of individual stimulus responses using pairwise category comparisons, enabling a more fine-grained and conservative assessment of category preference. For untrained ANNs, several model instances had no category-selective units exceeding this stringent threshold; thus, we adopted a more permissive cutoff of *t >* 3 (two-sided *p* = 2.82 × 10^−3^, *df* = 574) to obtain reliable category-selective unit sets across all architectures and seeds.

#### 4.2.3 Layer selection

To ensure a fair comparison between ANNs and the brain, we selected, for each fROI, the layer with the highest brain–model correspondence using a data-driven criterion. To do so, we treated all intermediate feature extraction layers as candidate layers, excluding the final classification (MLP) layers. In convolutional neural networks (CNNs), this included convolutional layers, normalization and nonlinearity stages, and the final pooling layer preceding the classifier. In Transformer-based models, we considered all encoder blocks following patch and positional embeddings, and within each block, we evaluated activations at multiple stages: after the first normalization, after self-attention (before residual addition), after the second normalization, and after the MLP block (before residual addition). To preserve spatial information, individual tokens were treated independently. This procedure yielded a set of candidate layers for each model.

For a given fROI (e.g., FFA), we first identified category-selective units for the fROI’s preferred category (e.g., face-selective) in every candidate layer as described above. We then computed responses of these units to the 485-image NSD training set and compared them to voxel-averaged fROI responses from the four training subjects. Specifically, we calculated the univariate Pearson correlation between unit-averaged model responses and voxel-averaged brain responses across stimuli (see Methods 4.3). Correlations were noise-corrected to account for within-subject variability across stimulus repetitions (see Methods 4.3.1). The representative layer was defined as the layer with the maximum median noise-corrected correlation across subjects. This procedure was performed independently for each fROI, using face-, body-, and scene-selective units for FFA, EBA, and PPA, respectively.

#### 4.2.4 Selecting best-performing ANN models

To compare how well different ANN models capture category-selective responses observed in the human brain, we ranked models based on their correspondence with fMRI responses. For each model, we computed a global score using the 485-image training set. Specifically, for each fROI (FFA, EBA, and PPA), we estimated the noise-corrected univariate correlation between voxel-averaged responses and unit-averaged responses from the corresponding representative layer (see Methods 4.3, 4.3.1), and took the median across the four training subjects (1, 2, 5, and 7). The global model score was defined as the median of these fROI-specific scores across FFA, EBA, and PPA. The model with the highest global score was treated as the best ANN model. The full ranked list of models is provided in **Supplementary Table 1**.

#### 4.2.5 Cross-validating category-selective ANN responses

To assess whether category-selective units generalize beyond the localizer stimuli, we evaluated their responses on a held-out image set. Specifically, we measured responses of these units from the selected layer to images from a separate stimulus set [24], which contained 80 images each of faces, scenes, bodies, and objects (320 images total). Responses were averaged across units and z-scored across images. To assess the robustness of these responses, we performed a one-sided Mann–Whitney U test comparing responses to the preferred category with each non-preferred category. We also visualized the resulting unit-averaged responses for each category.

### 4.3 Comparing category-selective population responses in brains and ANNs

To compare category-selective population responses between ANNs and the human brain, we used two complementary correlation-based metrics.

#### Univariate response tuning comparison

We compared response magnitudes across stimuli using the Pearson correlation between responses from two populations (*r*_1,2_). For brain data, responses were averaged across category-selective voxels, and for ANNs, across category-selective units.

To account for measurement noise within each system, we applied a noise-normalization procedure based on split-half reliability (see Methods 4.3.1). Specifically, the raw correlation between systems was divided by the square root of the product of their within-system reliabilities (*r*_1,1_ and *r*_2,2_), estimated from repeated presentations of the same images. This yields the noise-corrected Pearson correlation:

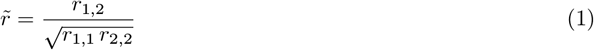

#### Multivariate response structure comparison

We compared the structure of population responses using representational similarity analysis (RSA) [48–50]. For each population, we constructed a response matrix across stimuli and voxels/units (*n*_stimuli_ × *n*_voxels/units_), and computed pairwise Euclidean distances between stimuli to obtain a representational dissimilarity matrix (RDM) (*n*_stimuli_ × *n*_stimuli_; see [54] for discussion of distance metrics).

We then quantified similarity between RDMs using Spearman’s rank correlation 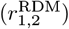. As above, we applied noise normalization using split-half reliability of the RDMs (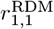 and 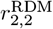), yielding the noise-corrected RDM correlation:

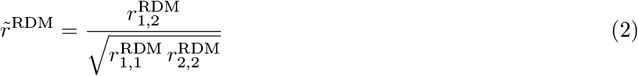

Note that because both 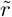and 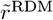 are ratios of empirically estimated correlations, they are not strictly bounded at 1. Values slightly above 1 can occur when the between-system correlation approaches the noise ceiling and sampling variability in the reliability estimates causes the denominator to underestimate the true within-system reliability. We report values without clipping to avoid introducing downward bias.

#### 4.3.1 Estimating split-half reliabilities

To estimate the reliability of responses within each system, we computed split-half reliability for both ANN units and fMRI voxels. Because ANN models are deterministic, their unit responses are noise-free, and split-half reliability was therefore set to 1.

For voxel responses from the NSD dataset, each image was presented three times. To estimate reliability, we used a split-half procedure in which, for each repetition *i* ∈{1, 2, 3}, responses from repetition *i* were correlated with the mean response across the other two repetitions (e.g., 1 vs. mean(2, 3), 2 vs. mean(1, 3), 3 vs. mean(1, 2)). Correlations were computed across images in the set (using voxel-averaged responses for univariate analyses and voxel-pattern correlations for multivariate analyses). Averaging across repetitions provides a more stable estimate of the underlying response pattern and reduces measurement noise.

Each raw split-half correlation was then Spearman–Brown corrected to estimate the reliability expected for the full dataset:

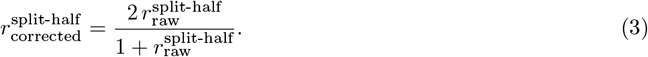

This procedure yielded three corrected reliability estimates for each population. The final reliability was defined as the maximum of these estimates to mitigate downward bias from noisy repetitions, which can be substantial with few repeats.

#### 4.3.2 Establishing the human-human ceiling

To estimate the human-human ceiling, we computed noise-corrected correlations across human subjects, following the NeuroAI training test [47]. For each subject, we calculated the median correlation with all other subjects. This procedure yields a distribution of human-human similarity, which serves as an upper bound on model performance. Ceilings were estimated independently for each metric (univariate and multivariate) and each fROI.

#### 4.3.3 Comparison of human-ANN similarity to the human-human ceiling

We tested whether ANN category-selective responses reached the human-human ceiling at both the individual model level and across the population of models.

At the individual model level, for each model and subject, we computed the difference between the human-ANN noise-corrected correlation and the subject-specific inter-subject ceiling (defined as the median noise-corrected correlation between that subject and all other subjects). To test whether human-ANN similarity fell below this ceiling, we performed an exact paired sign-flip permutation test across subjects (2^8^ permutations, one-sided), using the median difference across subjects as the test statistic.

At the population level, each model was summarized by its median difference across subjects. We then tested whether these model-level differences were below zero using a one-sided Wilcoxon signed-rank test across models (N = 35).

#### 4.3.4 Comparing human-human correlations across fROIs

To contextualize the gap between model–brain similarity and the human-human ceiling for a given fROI, we compared cross-subject correlations within and across fROIs. For each subject *s*, we computed noise-corrected correlations between that subject’s responses in fROI_1_ and the responses of every other subject *s*^*′*^ in both fROI_1_ and fROI_2_. For each comparison subject *s*^*′*^, we calculated the difference between the within-region correlation 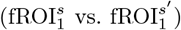 and the cross-region correlation 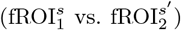. The gap for subject *s* was defined as the median of these differences across all comparison subjects. The final gap estimate was computed as the median of these subject-level gaps across subjects.

#### 4.3.5 Low-dimensional visualization of population response similarity

To visualize the similarity between category-selective responses in human brains and ANN models, we applied classical Multi-Dimensional Scaling (MDS). MDS embeds items in a low-dimensional space such that Euclidean distances approximate their pairwise dissimilarities. For each pair of populations (across all combinations of human subjects and ANN models), we computed noise-corrected similarity using either univariate correlations 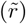 or multivariate RDM correlations 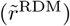, as described above. These similarities were converted to dissimilarities using

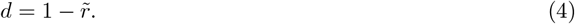

Dissimilarities were computed independently within each fROI (FFA, EBA, and PPA), and the final dissimilarity between each pair of populations was defined as the median across fROIs. Because noise-normalized correlations can occasionally produce negative distances due to variability in reliability estimates, we clipped all distances to a minimum value of 10^−5^ to ensure numerical stability. The resulting distance matrix (subjects and ANN models × subjects and ANN models) was embedded into a two-dimensional space using classical MDS (implemented via scikit-learn). The resulting embeddings reflect the relative dissimilarity structure of response patterns across all populations.

### 4.4 Investigating the effect of experimenter-defined parameters on category-selective response similarity between brains and ANNs

To assess how experimenter-defined choices in the analysis pipeline influence category-selective response similarity between brains and ANNs, we systematically varied key parameters, including the category-selectivity threshold and the functional localizer.

#### 4.4.1 Category-selectivity threshold

We examined how varying the category-contrast t-threshold for voxel and unit selection affected response similarity. Keeping the functional localizer fixed, we varied the threshold independently for brains and ANNs (using the previously selected representative layer) from *t >* 3 to *t >* 11 in increments of 2, yielding five threshold values. For each threshold, we quantified the number of selected voxels (in brains) and the proportion of selected units (in ANNs, relative to the total number of units in the layer), summarizing across subjects and models using the median. Split-half reliabilities for brain responses were re-estimated independently at each threshold. We then repeated the full analysis pipeline to compute category-selective response similarity between brains and ANNs across all pairs of thresholds (25 total pairs). To evaluate whether models reached the human-human ceiling, we applied the same statistical tests described above (see Methods 4.3.3). To visualize deviations from the ceiling, we computed a gap measure for each model, defined as the difference (median across subjects) between the model–brain correlation and the subject-specific human-human ceiling. We plotted the median gap across models for each threshold pair. As a supplementary analysis, we also report this gap for the best-performing model.

#### 4.4.2 Functional localizer

We next examined the effect of the functional localizer on category-selective unit identification. Keeping the threshold fixed (*t >* 7), we repeated the localization procedure in the previously selected representative layers of all ANN models using an alternative functional localizer [112]. This localizer consists of colored videos of faces, bodies, scenes, objects, and scrambled objects. We extracted frames from these videos to construct an image set, matching the number of images per category to the original localizer (144 scrambled; 288 per other category). To ensure diversity, we sampled five frames per video at uniform intervals (0%, 25%, 50%, 75%, and 100% of the video duration). Using this alternative localizer, we re-identified face-, scene-, and body-selective units. We then repeated the full analysis pipeline to compute category-selective response similarity between brains and ANNs using the alternative localizer.

To assess the stability of ANN category-selective unit identification across localizers, we quantified the overlap between unit sets identified using the two localizers via Intersection over Union (IoU), defined as the fraction of units shared across localizers relative to the union of identified units. We further established a ceiling for this identification procedure by measuring within-localizer consistency. Specifically, we computed a split-half IoU by dividing the stimulus set into two halves, repeating the identification procedure independently for each half, and computing the IoU between the resulting unit sets. This procedure was repeated over 50 random splits, and the final consistency estimate was taken as the median split-half IoU. Finally, to test whether across-localizer overlap was lower than within-localizer consistency, we performed a one-sided Wilcoxon signed-rank test across models for each category.

#### 4.4.3 Voxel-tuning matched ANN unit identification procedure

To test whether ANN unit subsets with more brain-like responses exist outside those identified by functional localizers, we performed a voxel–unit tuning matching analysis. For each ANN model and fROI, we considered units within the previously selected fROI-corresponding layer. Using the shared training image set (485 images), we computed the Pearson correlation between the stimulus tuning profiles of each voxel and each ANN unit across images. Correlations were noise-corrected using voxel split-half reliability (unit reliability was assumed to be 1). For each voxel, we selected the ANN unit whose responses were most strongly correlated with that voxel’s responses, yielding a one-to-one voxel–unit mapping (allowing a unit to be matched to multiple voxels). This procedure was performed independently for each subject (1, 2, 5, and 7), and the final unit subset for each model and fROI was defined as the union of units selected across subjects. We then repeated the full analysis pipeline using these matched unit subsets to estimate human-ANN response similarity and assessed whether human-ANN correlations reached the human-human ceiling using the same statistical tests described above (see Methods 4.3.3). As a control, we also evaluated a variant that enforced a unique unit assignment per voxel using a greedy matching procedure that iteratively selected the highest-correlation voxel–unit pair and removed both from further consideration. This constraint did not qualitatively affect the results.

#### 4.4.4 Choice of category-selective fROI

To assess the generality of our findings across category-selective regions, we repeated the analysis using an expanded set of fROIs in the NSD dataset, including OFA, FBA (FBA-1 and FBA-2), OPA, VWFA (VWFA-1 and VWFA-2), and OVWFA. For each fROI, voxels from both hemispheres were included. For ANNs, we used previously identified face-, body-, and scene-selective unit sets. Because word-selective units were not identified in the main analysis, we repeated the identification procedure to localize them. For each fROI, we re-identified the representative layer in all ANN models and then repeated the full analysis pipeline to estimate category-selective response similarity between brain responses and the corresponding ANN unit populations. Human-human ceilings and split-half reliabilities were estimated independently for each fROI. As before, statistical comparisons to the ceiling were performed using the same procedures described above (see Methods 4.3.3). We then tested whether model–brain correlations reached the human-human ceiling for each fROI (see Methods 4.3.3).

### 4.5 Voxel-wise encoding models

To test whether ANN unit populations can predict human brain responses, we modeled each voxel as a linear combination of ANN unit responses using an encoding model framework [45, 56–60, 85]. We considered three types of unit populations:

1. **All units**: using all the units in an ANN layer.
2. **Lesioned units**: using all units in the ANN layer except those previously localized as category-selective for the preferred category.
3. **Only category-selective units**: using only the previously localized category-selective units corresponding to the preferred category.

For each ANN model, unit populations were extracted from the previously selected layer corresponding to the target fROI. These features were used to predict voxel responses in the corresponding brain region (FFA, EBA, or PPA) for four subjects (1, 2, 5, and 7). Models were trained on the 485-image training set and evaluated on the held-out 515-image test set, as defined previously. All encoding models were fit and evaluated within each subject. For each voxel, we trained a ridge regression model (*α* = 0.1); further optimization of *α* did not meaningfully change the results. Prediction accuracy was quantified using univariate Pearson correlation and multivariate RDM correlation, as defined above. Because the evaluation was performed within-subject, noise correction was not applied. Instead, split-half reliability provided a within-subject ceiling for each fROI. For each model and unit population, predictive performance was summarized as the median prediction accuracy across subjects, with 95% confidence intervals. As supplementary analyses, we repeated the pipeline across different category-selectivity thresholds (*t >* 3 to *t >* 11) and using sparse-positive mappings via Lasso regression with non-negative weights (*α* = 0.01) [24, 62].

### 4.6 Interpreting the discrepancy between category-selective responses in brains and ANNs

To identify stimulus-level factors underlying discrepancies between category-selective responses in the brain and ANNs, we first constructed candidate feature groups based on systematic differences in response patterns.

For each fROI (FFA, EBA, and PPA), we examined univariate responses (voxel- or unit-averaged) from a representative subject (Subject 1) and a reference model (ResNet-50 ImageNet-1K) to the shared evaluation set (515 images). We identified groups of stimuli that elicited divergent responses between the two systems and manually extracted common visual or semantic features across these groups to generate hypotheses about the sources of these divergences.

We evaluated these hypotheses using held-out images, subjects, and models. Specifically, for each group, we selected 25 images per subject from subject-specific stimulus sets (approximately 9000 images per subject), ensuring no overlap with training or evaluation images. Only stimuli with at least one repetition were included. For each subject, responses were z-scored across the full subject-specific stimulus pool, and this normalization was applied independently to brain and ANN responses to express both relative to the same stimulus distribution. We evaluated generalization across subjects (7 held-out subjects) and models (top 25% of models based on the global score; 10 models; see **Supplementary Table 1**).

To quantify differences between brain and ANN responses, we fit a linear mixed-effects model for each preferred category and model. The response for trial *i* was modeled as

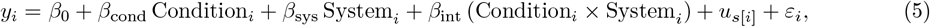

where Condition_*i*_ denotes the stimulus group (Group 1 vs. Group 2) and System_*i*_ denotes the response source (brain vs. ANN), with reference levels Group 2 and ANN. The term *u*_*s*[*i*]_ ∼ 𝒩 (0, *σ*^2^) represents a subject-specific random intercept, and *ε*_*i*_ ∼ 𝒩 (0, *σ*^2^) denotes residual error. The Condition × System interaction (*β*_int_) was used to assess divergence between brain and model responses. Statistical inference was based on Wald *z*-tests with one-sided *p*-values for directional hypotheses.

## Acknowledgements

This work was supported by a CoCo fellowship by the School of Psychology, Georgia Tech, to A.D.; the NIH Pathway to Independence Award (R00EY032603), NSF Nexus (Allocation number: SOC250049), and a startup grant from Georgia Tech to N.A.R.M. We are grateful to Doby Rahnev, Anya Ivanova, and Micheal Cohen, and members of the Murty Lab for feedback on the manuscript. We would like to thank the members of Murty Lab, Ivanova Lab, Rahnev Lab, Saloni Sharma, and Kasper Vinken for many thoughtful discussions and feedback on the paper.

## Author contributions

A.D. and N.A.R.M. conceived and planned the research; A.D. and N.A.R.M. performed research; A.D. analyzed data; and A.D. and N.A.R.M. wrote the paper.

## Data and materials availability

Data and code will be made publicly available upon acceptance.

## 5 Supplemental Information

### Supplemental Figures

**S1**. Trained ANN models have robust face-, body- and scene-selective units

**S2**. Untrained ANN models do not have robust face-, body- and scene-selective units

**S3**. Noise-corrected univariate correlations across all subjects and models, for all category-selective regions

**S4**. Noise-corrected multivariate correlations across all subjects and models, for all category-selective regions

**S5**. Univariate and multivariate correlation gaps between brains and best-performing model across combinations of brain and ANN selectivity thresholds

**S6**. Intersection of unit sets identified within and across different localizers

**S7**. Noise-corrected univariate and multivariate correlations for voxel-matched ANN unit subsets

**S8**. Noise-corrected univariate and multivariate correlations across an extended set of category-selective regions.

**S9**. Prediction accuracy of voxel-wise encoding models when category-selective units are lesioned, across selectivity thresholds

**S10**. Prediction accuracy of voxel-wise encoding models when category-selective units are lesioned, using sparse-positive mapping

**S11**. Brain and ANN responses for chosen stimulus groups, across all selectivities

**S12**. Brain and ANN responses for extended stimulus groups, across all selectivities

**S13**. Brain and ANN responses for all stimulus groups, including random, across all selectivities

### Supplemental Tables

**S1**. Summary of all the pretrained models evaluated

**S2**. Summary of all the untrained model architectures evaluated

---

**Figure S1.**
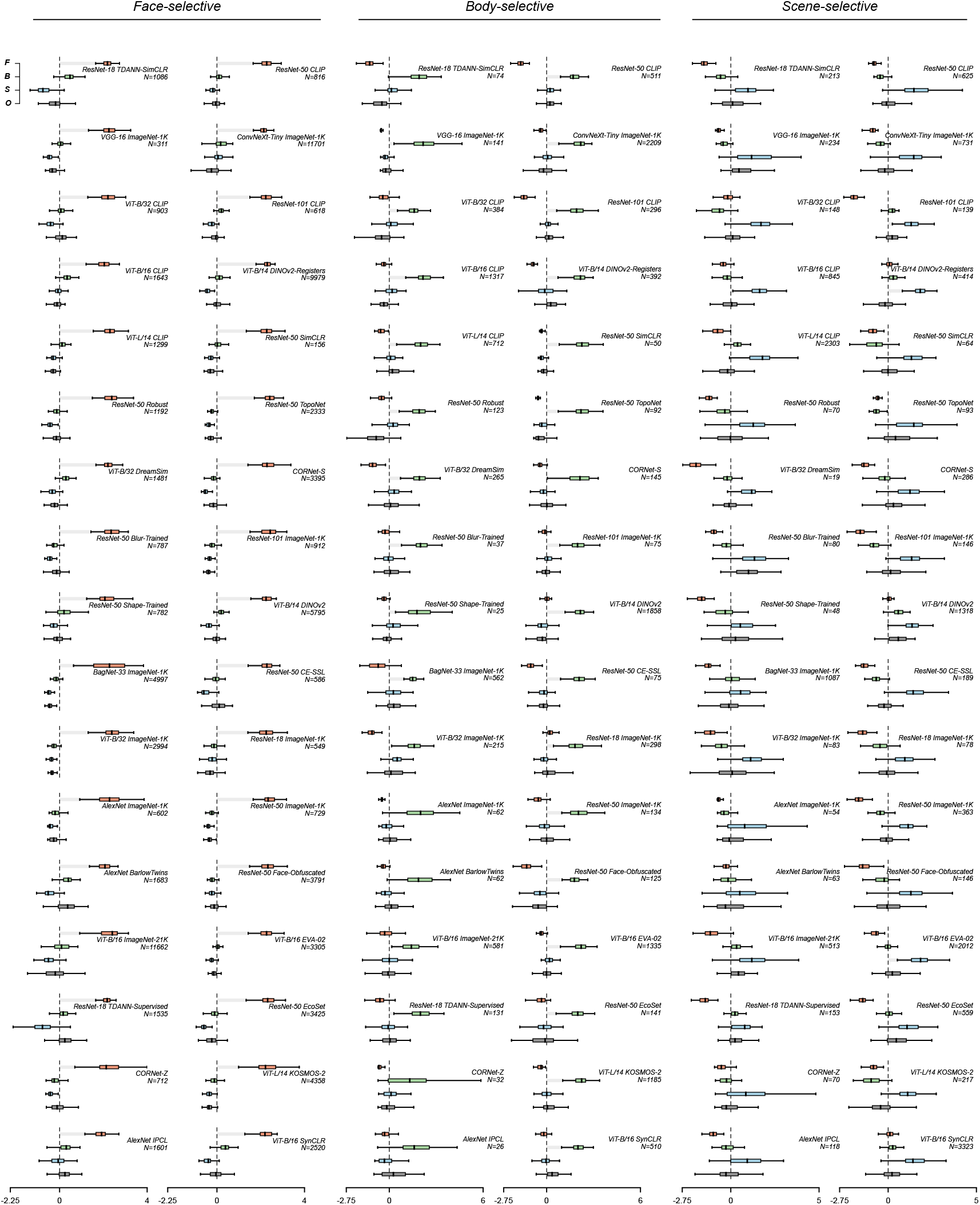
Trained ANN models have robust face-, body- and scene-selective units. Responses of localized face-, body-, and scene-selective units to independent stimuli of faces (F), bodies (B), scenes (S), and objects (O). For all boxplots, the y-axis indicates the stimulus category, and the x-axis shows the unit-averaged z-scored responses for a model. The annotated text shows the model name and the number of units localized. The grouped columns show responses of face-, body-, and scene-selective units, respectively. The best-performing model (ViT-B/16 SigLIP2) is repeated from Fig. 2.

**Figure S2.**
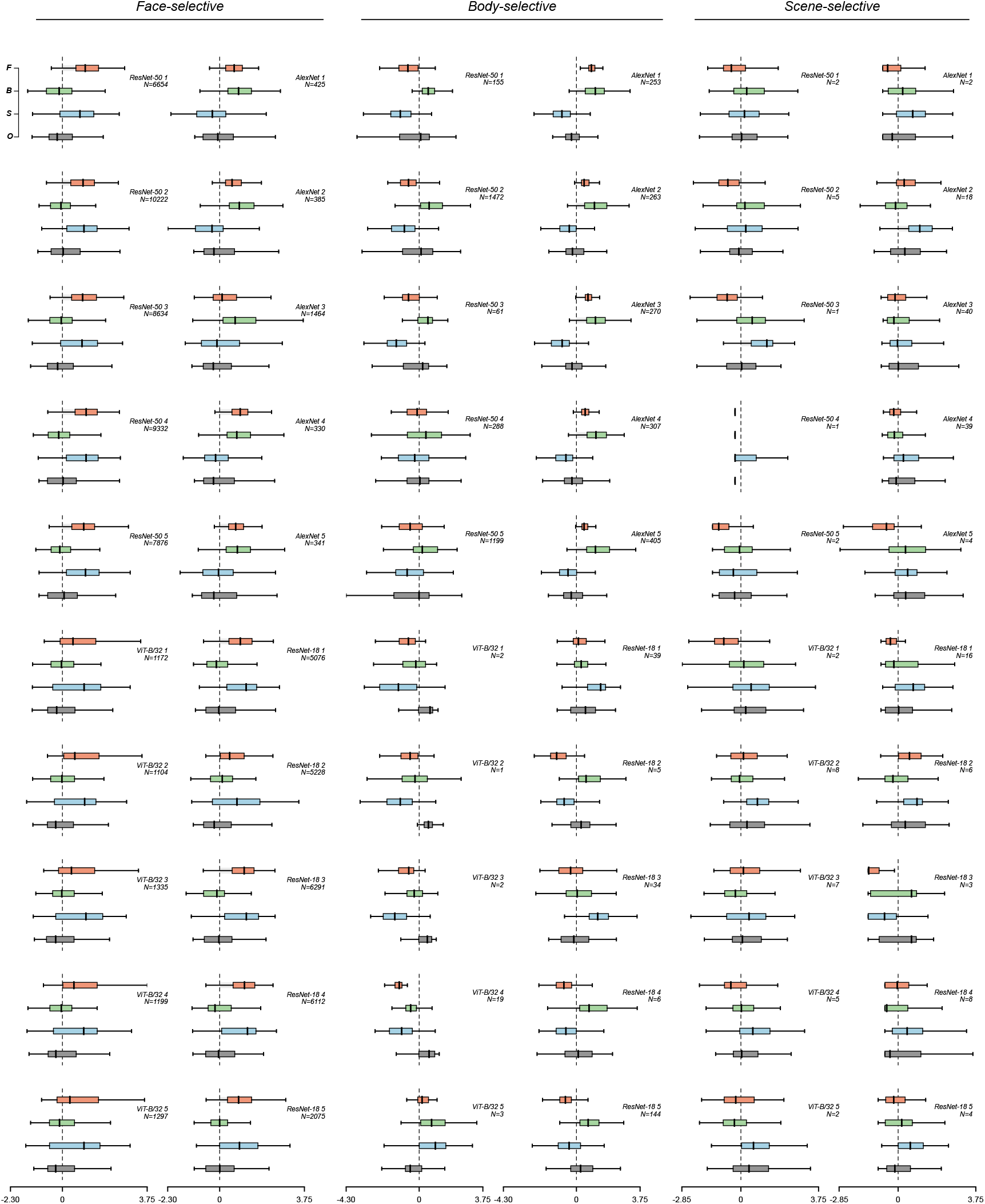
Untrained ANN models do not have robust face-, body- and scene-selective units. Responses of localized face-, body-, and scene-selective units to independent stimuli of faces (F), bodies (B), scenes (S), and objects (O). For all boxplots, the y-axis indicates the stimulus category, and the x-axis shows the unit-averaged z-scored responses for a model. The annotated text shows the model name and the number of units localized. The grouped columns show responses of face-, body-, and scene-selective units, respectively. AlexNet 5 is repeated from Fig. 2.

**Figure S3.**
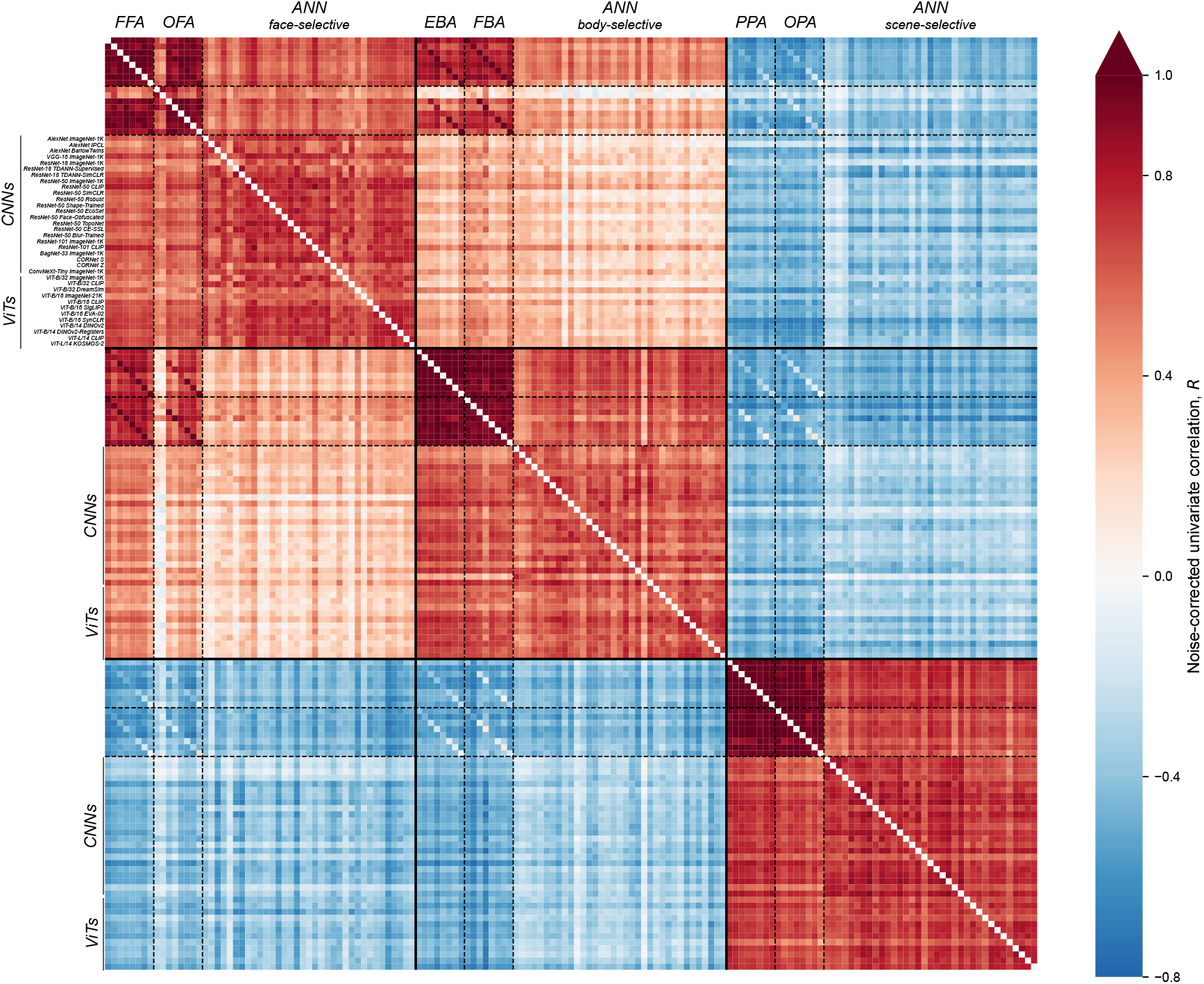
Noise-corrected univariate correlations across all subjects and trained models, for all category-selective regions. Noise-corrected univariate correlations between all pairs of subjects and models, across all selectivities and an extended set of category-selective regions. ANN representative layers correspond to canonical category-selective regions, i.e., FFA, EBA, and PPA. Subjects are arranged numerically (1-8), and models are grouped based on their architectures, with model names annotated.

**Figure S4.**
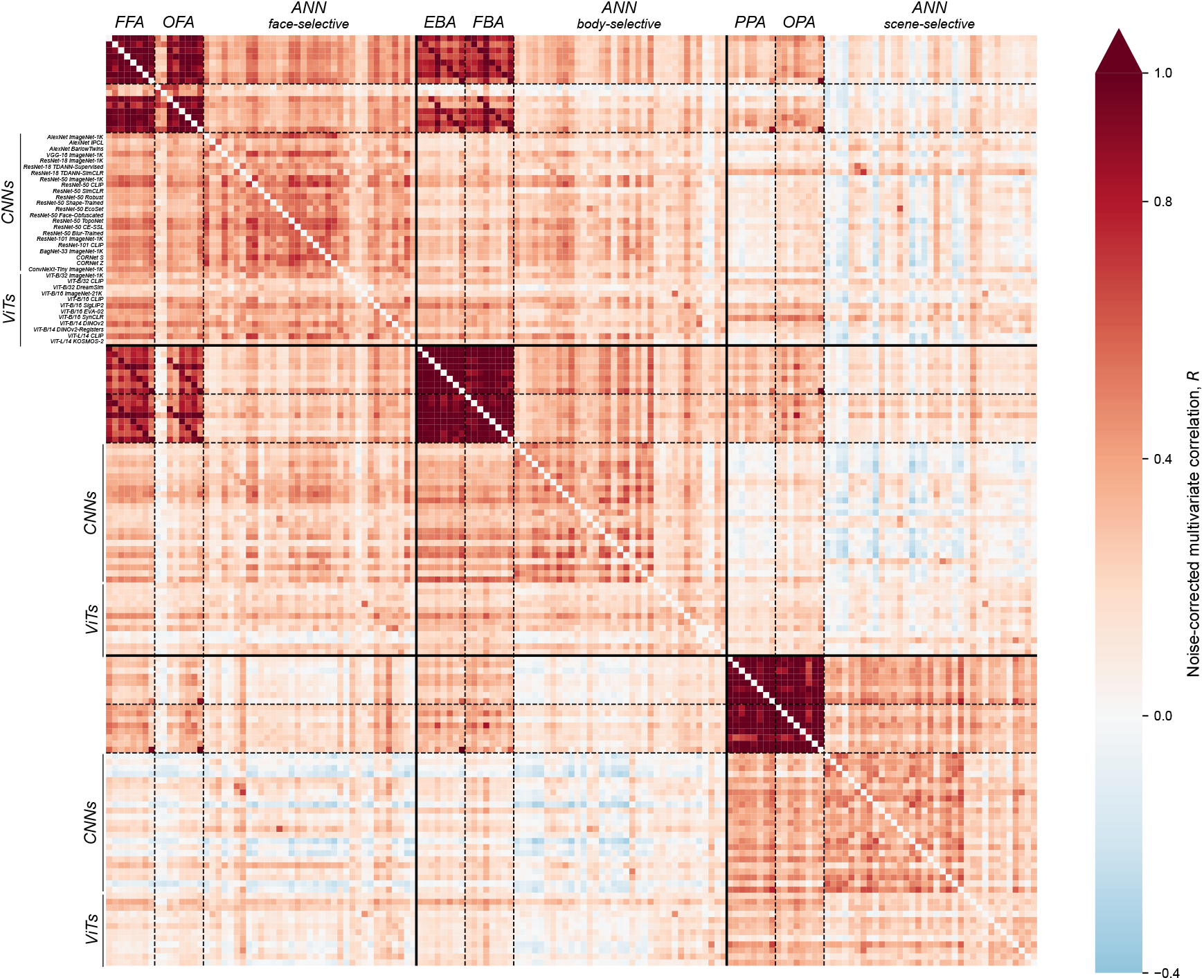
Noise-corrected multivariate correlations across all subjects and trained models, for all category-selective regions. Noise-corrected multivariate correlations between all pairs of subjects and models, across all selectivities and an extended set of category-selective regions. ANN representative layers correspond to canonical category-selective regions, i.e., FFA, EBA, and PPA. Subjects are arranged numerically (1-8), and models are grouped based on their architectures, with model names annotated.

**Figure S5.**
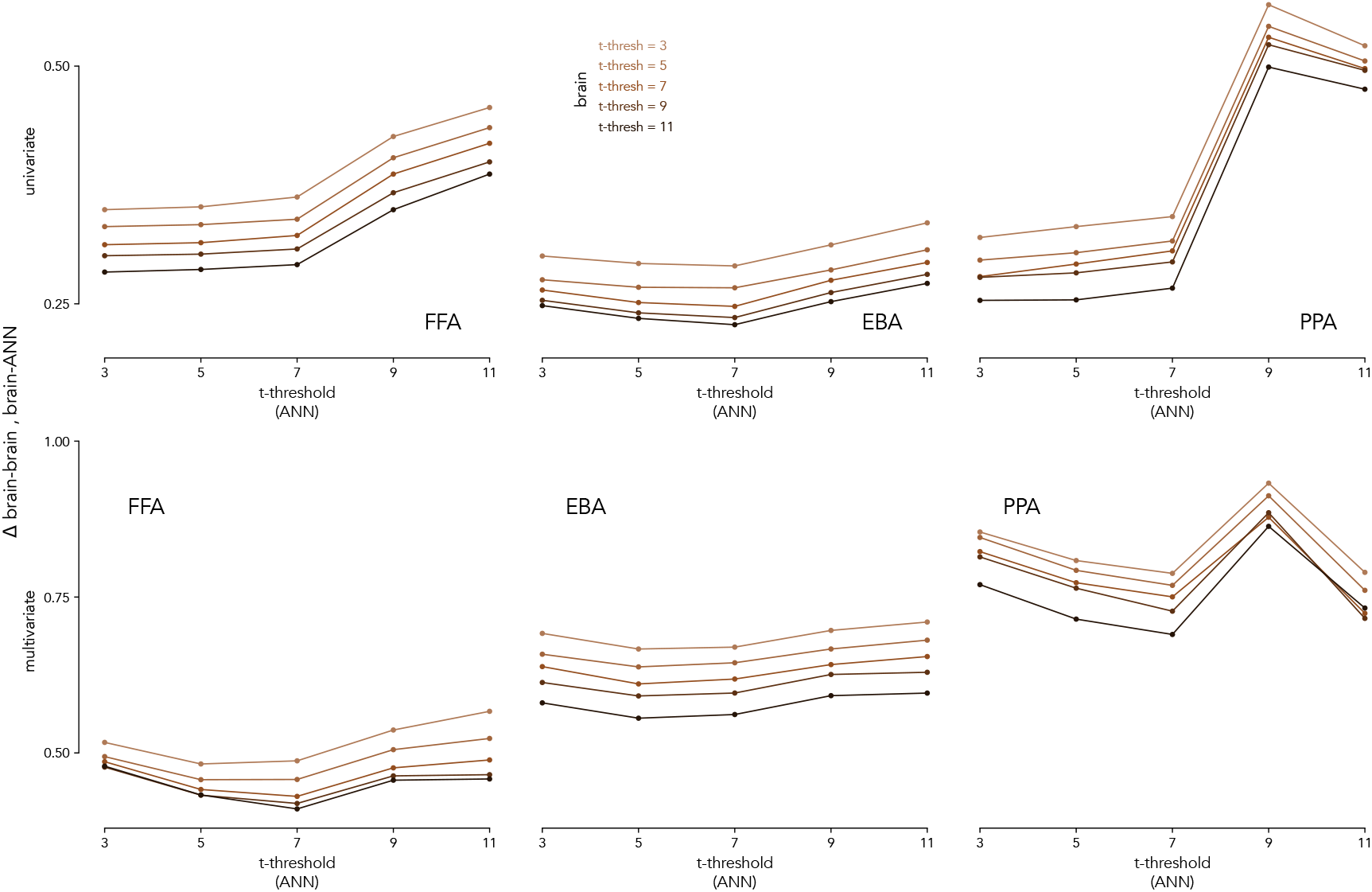
Univariate and multivariate correlation gaps between brains and best-performing model across combinations of brain and ANN selectivity thresholds. Line plots show the noise-corrected correlation gap between the best model (ViT-B/16 SigLIP2) and brains across category-selective regions and comparisons. For each region, the x-axis shows the ANN t-threshold, whereas the y-axis shows the gap in noise-corrected correlations. The gap is the difference between a subject’s median correlation with other subjects and correlation with the model (median across subjects). Lines of different shades show the gap for sets of voxels corresponding to different values of Brain’s t-threshold, with darker shades corresponding to higher t-threshold. The top row shows the correlation estimated using univariate comparisons, and the bottom row shows the correlations estimated using multivariate comparisons.

**Figure S6.**
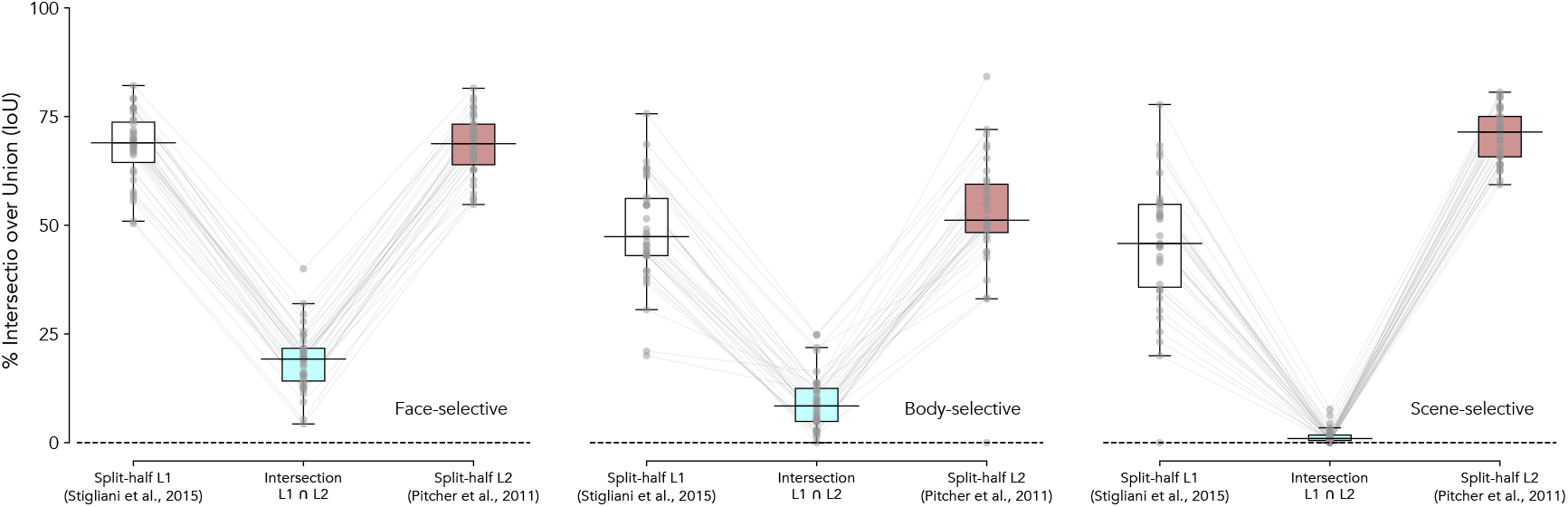
Intersection of unit sets identified within and across different localizers. Boxplots show the percentage overlap (Intersection over Union; IoU) between unit subsets identified using two different functional localizers (intersection *L*1 ∩ *L*2; same as Fig. 4), compared with the within-localizer split-half consistency for each localizer (*L*1 and *L*2). Each point represents one ANN model, and lines connect values from the same model. Across all selectivity domains (face, body, and scene), the overlap between units identified by different localizers is substantially lower than the within-localizer split-half overlap, indicating that the specific stimulus set used for localization strongly influences which ANN units are identified as category-selective.

**Figure S7.**
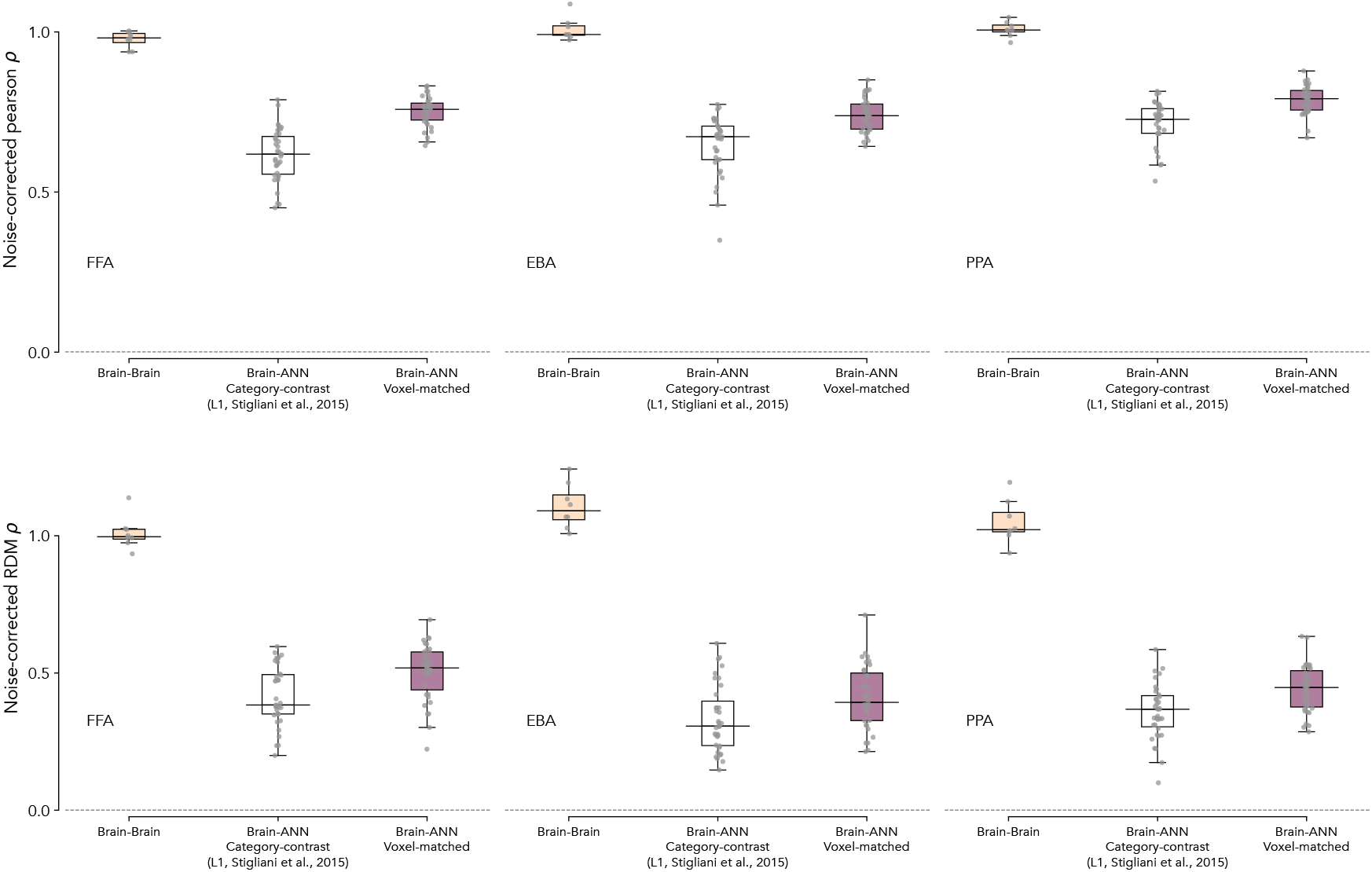
Noise-corrected univariate and multivariate correlations for voxel-matched ANN unit subsets. Descriptive boxplots illustrating the category-selective region (x-axis) v/s. noise-corrected correlation (y-axis) when evaluated on the NSD dataset. For each region, the first boxplot depicts the correlation of responses between human brains (each point indicates one subject’s median correlation with all other subjects); the second and third boxplots depict the correlation of responses between the ANN models and human brains for category-selective units identified using functional localizer 1 (same as Fig. 2b) and through voxel-tuning matching procedure, respectively (each point indicates one ANN model’s median correlation across all subjects). The ANN representative layer corresponding to fROI is the same as selected previously using functional localizer 1. The top row shows the correlation estimated using univariate comparisons, and the bottom row shows the correlations estimated using multivariate comparisons.

**Figure S8.**
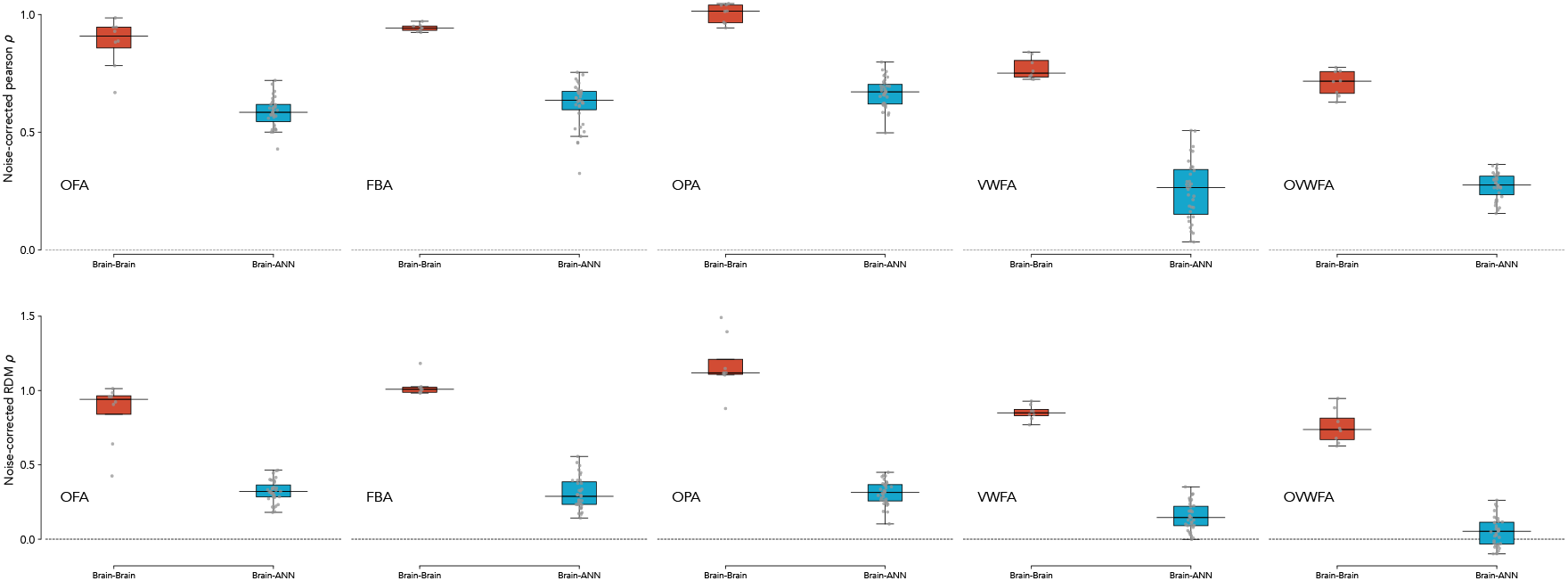
Noise-corrected univariate and multivariate correlations across an extended set of category-selective regions. Descriptive boxplots illustrating the category-selective region (x-axis) v/s. noise-corrected correlation (y-axis) when evaluated on the NSD dataset. For each region, the left boxplot depicts the correlation of responses between human brains (each point indicates one subject’s median correlation with all other subjects); the right boxplot depicts the correlation of responses between the ANN models and human brains (each point indicates one ANN model’s median correlation across all subjects). Here, the category-selective units were localized using functional localizer 1 (same as Fig. 2), and the layer was selected independently for each fROI. The top row shows the correlation estimated using univariate comparisons, and the bottom row shows the correlations estimated using multivariate comparisons.

**Figure S9.**
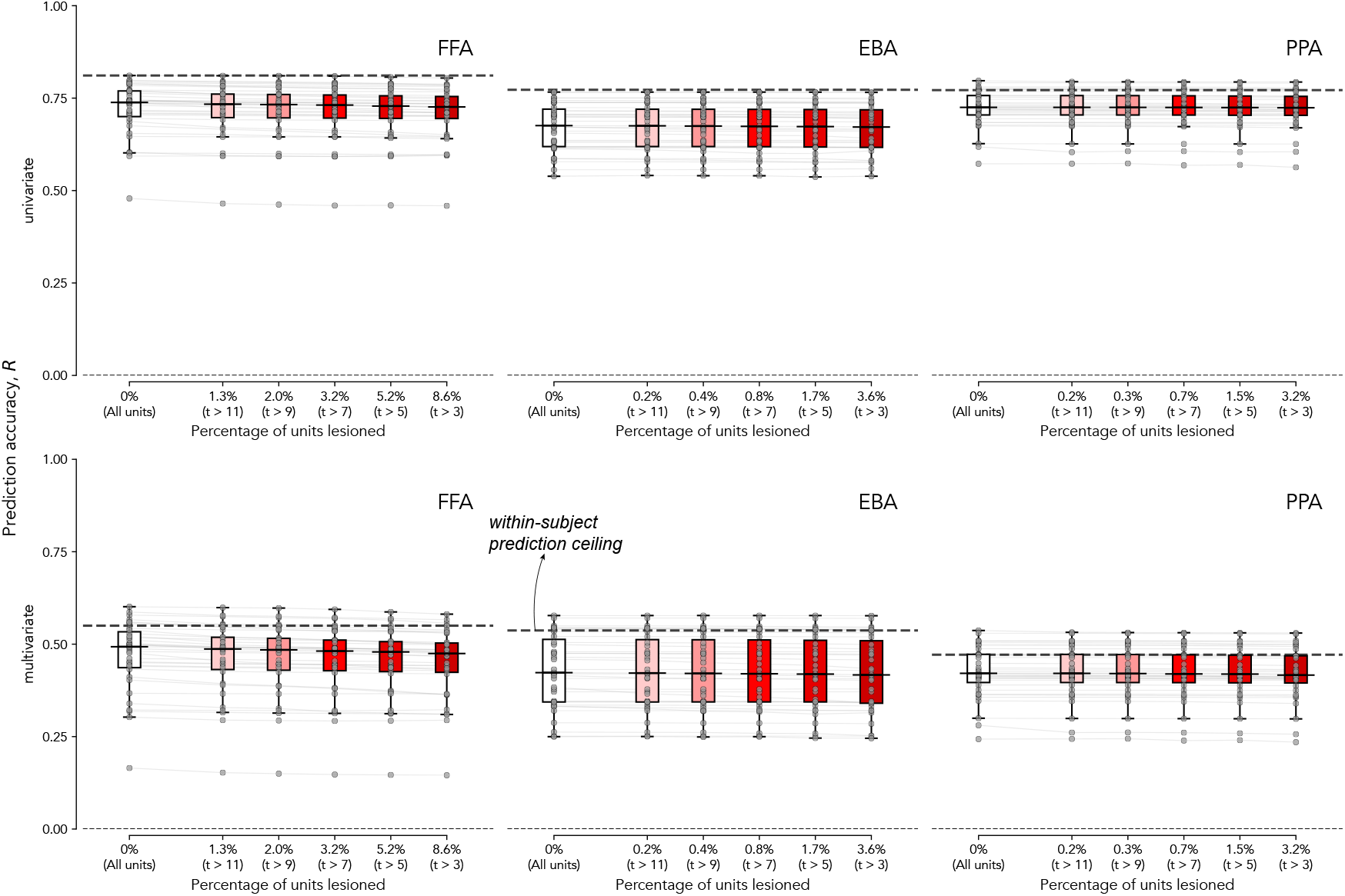
Prediction accuracy of voxel-wise encoding models when category-selective units are lesioned, across selectivity thresholds. Correlation between observed and predicted responses from voxel-wise encoding models (using ridge mapping) on held-out NSD images. For each boxplot, the x-axis indicates the percentage of category-selective units lesioned (category corresponding to the fROI’s preferred category), and the y-axis indicates the correlation. The x-axis labels also annotate the corresponding selectivity threshold for the category-selective unit identification. Each box depicts the correlation of the encoding model (each point indicating one ANN model’s median correlation across the subjects). The first box indicates the baseline using all units in the layer (previously chosen representative layer corresponding to the fROI). The black dashed line shows the median within-subject ceiling, i.e., spearman-brown corrected split-half correlation between the subject’s responses during the three repetitions of the same stimuli. The two rows indicate univariate and multivariate comparisons, respectively. The three columns indicate the category-selective region, i.e., FFA, EBA, and PPA, respectively.

**Figure S10.**
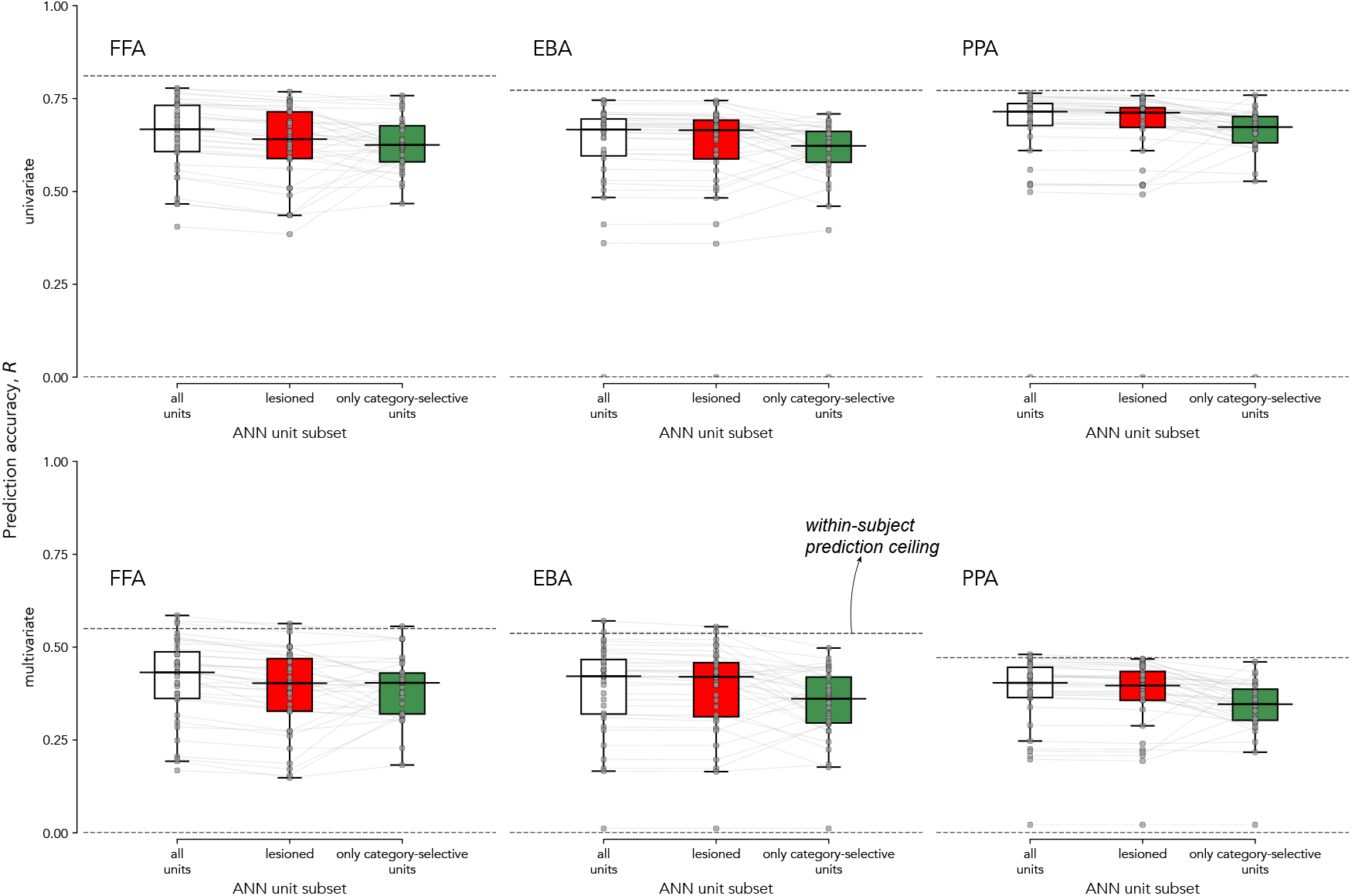
Prediction accuracy of voxel-wise encoding models when category-selective units are lesioned, using sparse-positive mapping. Correlation between observed and predicted responses from voxel-wise encoding models (using sparse-positive mapping) on held-out NSD images. For each boxplot, the x-axis indicates the unit set considered, and the y-axis indicates the correlation. The first box depicts the correlation of the encoding model using all the units within the ANN layer (each point indicating one ANN model’s median correlation across the subjects); similarly, the second and third boxes correspond to encoding models when the category-selective units are lesioned, and when only category-selective units are considered. The black dashed line shows the median within-subject ceiling, i.e., spearman-brown corrected split-half correlation between the subject’s responses during the three repetitions of the same stimuli. The two rows indicate univariate and multivariate comparisons, respectively. The three columns indicate the category-selective region, i.e., FFA, EBA, and PPA, respectively.

**Figure S11.**
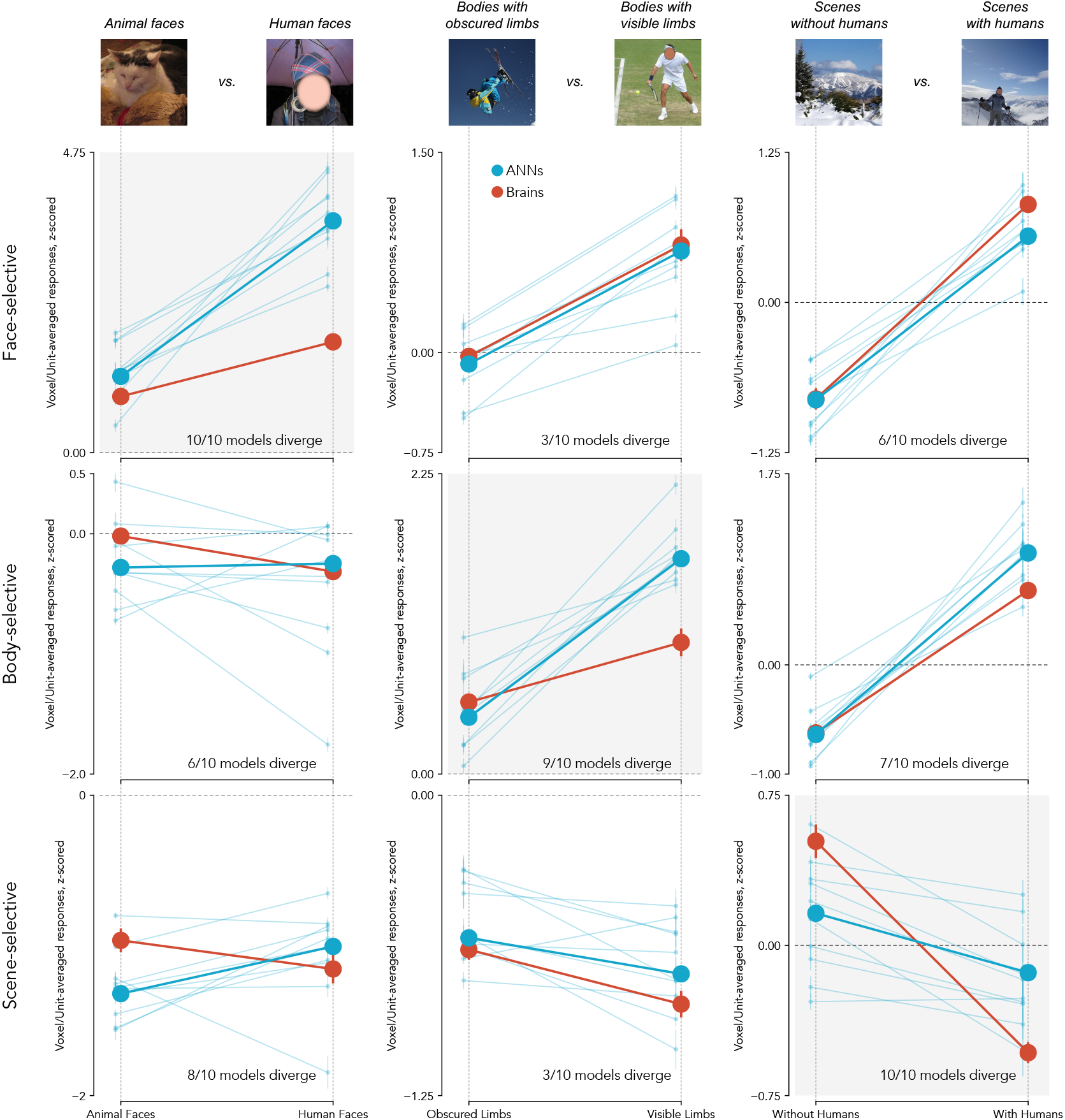
Brain and ANN responses for chosen stimulus groups, across all selectivities. Hypothesis validation of the candidate stimulus-level factors when evaluated on independent, subject-specific held-out images, held-out subjects, and held-out top-10 most brain-aligned models according to univariate tests. The top panel shows example images from the stimulus groups that were hypothesized to produce systematic divergence between ANN and brain responses from three domains of category selectivity: face-selective (animal faces vs. human faces), body-selective (bodies with obscured limbs vs. visible limbs), and scene-selective (scenes without humans vs. scenes with humans) (same as Fig. 6). A total of 350 stimuli were used for evaluation (25 per group per subject). The line plots show a descriptive visualization of ANN–brain divergence for each stimulus group. Three columns show the stimulus groups, and the three rows show the evaluated selectivities (and the corresponding fROI, i.e., FFA, EBA, and PPA, respectively). The shaded plot depicts hypotheses generated and evaluated within selectivity (here, the plots are the same as Fig. 6). For each plot, the x-axis indicates stimulus group, and the y-axis shows unit-averaged (ANN) or voxel-averaged (brain) z-scored responses. In all plots, the thick red lines denote the mean response across subjects (points indicate mean *±*SEM), thick blue lines denote the median response across the models, and faint blue lines show individual model responses. The annotated text shows the number of models (out of the top-10) that diverge, i.e., have a significant group × system interaction effect using the linear mixed-effects model as before.

**Figure S12.**
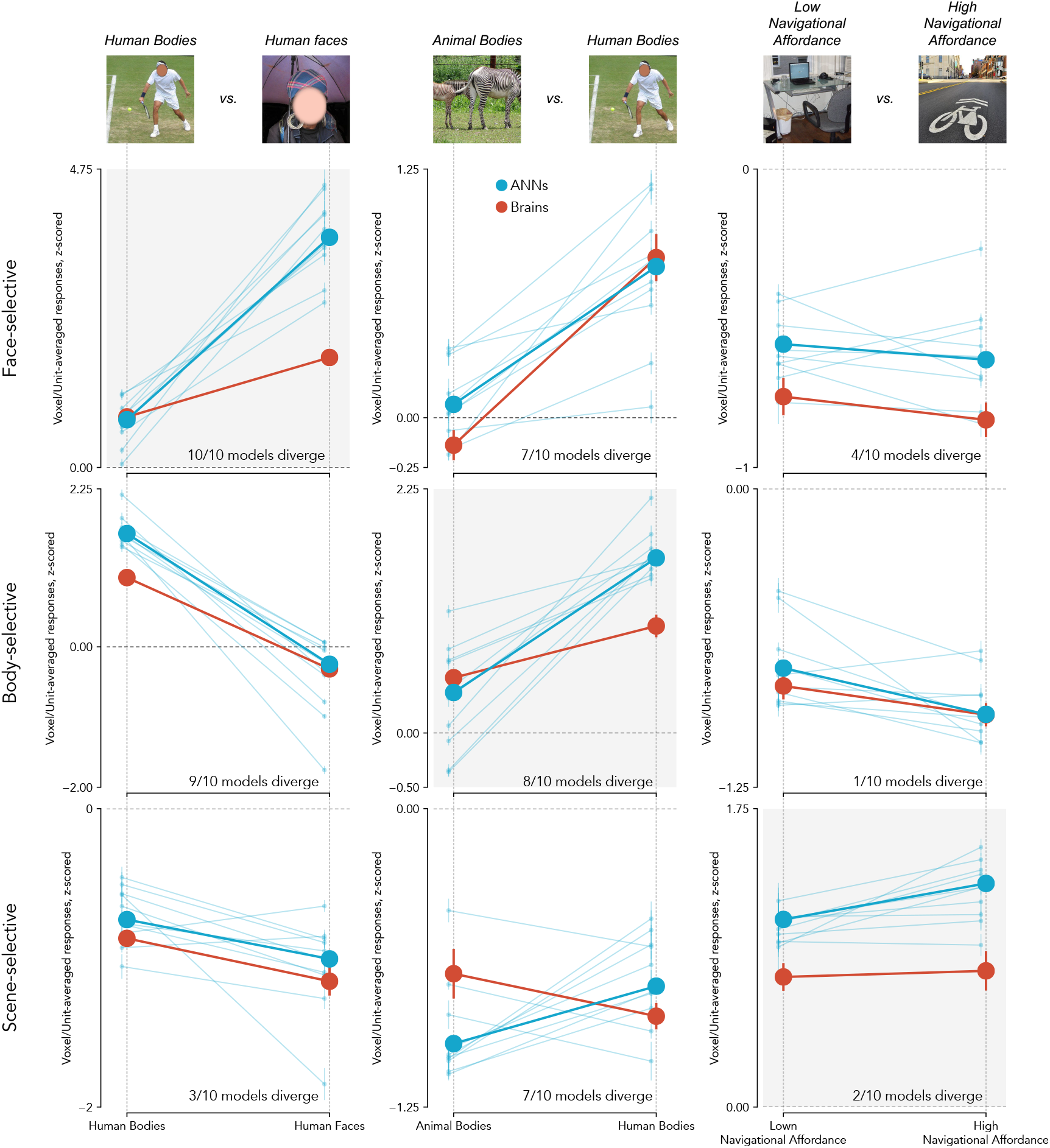
Brain and ANN responses for extended stimulus groups, across all selectivities. Hypothesis validation of the candidate stimulus-level factors when evaluated on independent, subject-specific held-out images, held-out subjects, and held-out top-10 most brain-aligned models according to univariate tests. The top panel shows example images from the stimulus groups that were hypothesized to produce systematic divergence between ANN and brain responses from three domains of category selectivity: face-selective (human bodies vs. faces), body-selective (animal vs. human bodies), and scene-selective (low vs. high navigational affordances) (human faces, and human bodies groups are the same as Fig. 6 and Fig. S11). A total of 350 stimuli were used for evaluation (25 per group per subject). The line plots show a descriptive visualization of ANN–brain divergence for each stimulus group. Three columns show the stimulus groups, and the three rows show the evaluated selectivities (and the corresponding fROI, i.e., FFA, EBA, and PPA, respectively). The shaded plot depicts hypotheses generated and evaluated within selectivity. For each plot, the x-axis indicates stimulus group, and the y-axis shows unit-averaged (ANN) or voxel-averaged (brain) z-scored responses. In all plots, the thick red lines denote the mean response across subjects (points indicate mean *±* SEM), thick blue lines denote the median response across the models, and faint blue lines show individual model responses. The annotated text shows the number of models (out of the top-10) that diverge, i.e., have a significant group × system interaction effect using the linear mixed-effects model as before.

**Figure S13.**
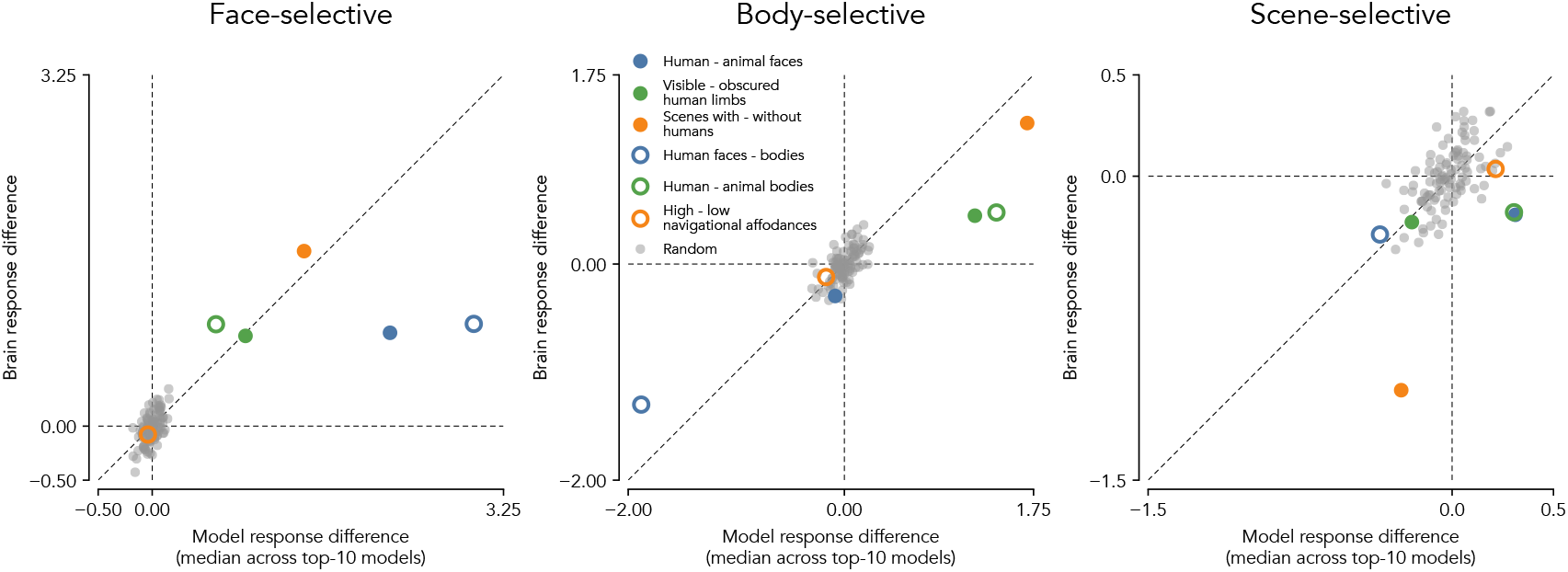
Brain and ANN responses for all stimulus groups, including random, across all selectivities. Each panel shows the difference in responses between the two stimulus groups for the brain (y-axis) and ANN models (x-axis). Brain response differences correspond to voxel-averaged responses in the corresponding category-selective region (FFA, EBA, or PPA) (median of within-subject difference, subjects 2-7), and model response differences correspond to unit-averaged responses from the top-10 most brain-aligned models (median of within-subject difference for each model, then median across models). Each colored point represents a stimulus contrast hypothesized to produce divergent responses between ANNs and brains across three domains of category selectivity: face-selective (animal faces vs. human faces), body-selective (bodies with obscured limbs vs. visible limbs), and scene-selective (scenes without humans vs. scenes with humans) (same as Fig. 6, S11). Additional contrasts tested across domains: face-selective (human bodies vs. faces), body-selective (animal vs. human bodies), and scene-selective (low vs. high navigational affordances) (same as Fig. S12). Gray points indicate control analyses using randomly sampled stimulus groups (100 bootstrap samples; each sample contains 350 stimuli total, 25 per group per subject) drawn from the same held-out subject-specific image sets (excluding the stimuli selected in previous hypotheses). These random contrasts cluster around the origin and do not show systematic divergence. The dashed diagonal line indicates equality between model and brain response differences. Points that lie far from this line indicate stimulus contrasts where ANN and brain responses diverge. Across selectivities, the hypothesized stimulus groups consistently show larger response differences in ANNs than in the corresponding brain regions, whereas random stimulus groups show no such divergence.

**Table 1.** Summary of all the pretrained models evaluated.

| # | Model name | Optimization Type | Architecture | Training Objective | Citation |
| --- | --- | --- | --- | --- | --- |
| 1 | ViT-B/16 SigLIP2 | Task-optimized | Transformer | Language-aligned | [113] |
| 2 | ResNet-50 CLIP | Task-optimized | CNN | Language-aligned | [109] |
| 3 | ConvNeXt-Tiny ImageNet-1K | Task-optimized | CNN | Supervised | [114] |
| 4 | ResNet-101 CLIP | Task-optimized | CNN | Language-aligned | [109] |
| 5 | ViT-B/14 DINOv2-Registers | Task-optimized | Transformer | Self-supervised | [115] |
| 6 | ResNet-50 SimCLR | Task-optimized | CNN | Self-supervised | [116] |
| 7 | ResNet-50 TopoNet<br>($\tau = 30$) | Task-optimized | CNN | Supervised | [29] |
| 8 | CORNet S | Task-optimized | CNN | Supervised | [96] |
| 9 | ResNet-101 ImageNet-1K | Task-optimized | CNN | Supervised | [117] |
| 10 | ViT-B/14 DINOv2 | Task-optimized | Transformer | Self-supervised | [118] |
| 11 | ResNet-50 CE-SSL<br>(BarlowTwins) | Task-optimized | CNN | Self-supervised | [119] |
| 12 | ResNet-18 ImageNet-1K | Task-optimized | CNN | Supervised | [117] |
| 13 | ResNet-50 ImageNet-1K | Task-optimized | CNN | Supervised | [117] |
| 14 | ResNet-50 Face-Obfuscated | Task-optimized | CNN | Supervised | [120] |
| 15 | ViT-B/16 EVA-02 | Task-optimized | Transformer | Language-aligned | [121] |
| 16 | ResNet-50 EcoSet | Task-optimized | CNN | Supervised | [92] |
| 17 | ViT-L/14 KOSMOS-2 | Task-optimized | Transformer | Language-Aligned | [122] |
| 18 | ViT-B/16 SynCLR | Task-optimized | Transformer | Self-supervised | [123] |
| 19 | ResNet-18 TDANN-SimCLR | Task-optimized | CNN | Self-supervised | [30] |
| 20 | VGG-16 ImageNet-1K | Task-optimized | CNN | Supervised | [124] |
| 21 | ViT-B/32 CLIP | Task-optimized | Transformer | Language-aligned | [109] |
| 22 | ViT-B/16 CLIP | Task-optimized | Transformer | Language-aligned | [109] |
| 23 | ViT-L/14 CLIP | Task-optimized | Transformer | Language-Aligned | [109] |

Table 1 – continued from previous page
| # | Model name | Optimization Type | Architecture | Training Objective | Citation |
| --- | --- | --- | --- | --- | --- |
| 24 | ResNet-50 Robust<br>(L2 $\epsilon = 3$) | Task-optimized | CNN | Supervised | [125] |
| 25 | ViT-B/32 DreamSim<br>(Open CLIP) | Task-optimized | Transformer | Language-Aligned | [126] |
| 26 | ResNet-50 Blur-Trained<br>(Strong) | Task-optimized | CNN | Supervised | [88] |
| 27 | ResNet-50 Shape-Trained | Task-optimized | CNN | Supervised | [127] |
| 28 | BagNet-33 ImageNet-1K | Task-optimized | CNN | Supervised | [128] |
| 29 | ViT-B/32 ImageNet-1K | Task-optimized | Transformer | Supervised | [129] |
| 30 | AlexNet ImageNet-1K | Task-optimized | CNN | Supervised | [130] |
| 31 | AlexNet BarlowTwins | Task-optimized | CNN | Self-supervised | [24, 131] |
| 32 | ViT-B/16 ImageNet-21K | Task-optimized | Transformer | Supervised | [132] |
| 33 | ResNet-18 TDANN-Supervised | Task-optimized | CNN | Supervised | [30] |
| 34 | CORNet Z | Task-optimized | CNN | Supervised | [96] |
| 35 | AlexNet IPCL<br>(alexnetgn_ipcl_ref01) | Task-optimized | CNN | Self-supervised | [110] |

**Table 2.** Summary of all the untrained model architectures evaluated.

| # | Model name | Optimization Type | Architecture | Training Objective | Citation |
| --- | --- | --- | --- | --- | --- |
| 1 | AlexNet | Untrained | CNN | - | [130] |
| 2 | ResNet-18 | Untrained | CNN | - | [117] |
| 3 | ResNet-50 | Untrained | CNN | - | [117] |
| 4 | ViT-B/32 | Untrained | Transformer | - | [129] |

